# Signal processing in auditory cortex underlies degraded speech sound discrimination in noise

**DOI:** 10.1101/833558

**Authors:** Stephen M. Town, Katherine C. Wood, Jennifer K. Bizley

## Abstract

The ability to recognize sounds in noise is a key part of hearing, and the mechanisms by which the brain identifies sounds in noise are of considerable interest to scientists, clinicians and engineers. Yet we know little about the necessity of regions such as auditory cortex for hearing in noise, or how cortical processing of sounds is adversely affected by noise. Here we used reversible cortical inactivation and extracellular electrophysiology in ferrets performing a vowel discrimination task to identify and understand the causal contribution of auditory cortex to hearing in noise. Cortical inactivation by cooling impaired task performance in noisy but not clean conditions, while responses of auditory cortical neurons were less informative about vowel identity in noise. Simulations mimicking cortical inactivation indicated that effects of inactivation were related to the loss of information about sounds represented across neural populations. The addition of noise to target sounds drove spiking activity in auditory cortex and recruitment of additional neural populations that were linked to degraded behavioral performance. To suppress noise-related activity, we used continuous exposure to background noise to adapt the auditory system and recover behavioral performance in both ferrets and humans. Inactivation by cooling revealed that the benefits of continuous exposure were not cortically dependent. Together our results highlight the importance of auditory cortex in sound discrimination in noise and the underlying mechanisms through which noise-related activity and adaptation shape hearing.

## Introduction

Signal processing in noise is a central requirement of many sensory systems. In hearing, everyday sounds are often experienced in noisy environments that present significant challenges for tasks such as speech and music perception. This is true both in healthy listeners and particularly for individuals with hearing loss. How the brain solves these challenges, or fails to do so, is thus an important question in auditory neuroscience

Many studies have asked how neurons encode sounds in noise at various stages of the auditory system (Nagarajan, Cheung et al. 2002, Bar-Yosef and Nelken 2007, Narayan, Best et al. 2007, Ding and Simon 2013, Teschner, Seybold et al. 2016, Ni, Bender et al. 2017) and emphasized the importance of auditory cortex in building noise-tolerant representations of sound identity. In primary and non-primary auditory cortex, neural responses to sounds in noise more closely match responses to corresponding clean sounds than the original degraded stimuli (Moore, Lee et al. 2013, Mesgarani, David et al. 2014, Fuglsang, Dau et al. 2017, Kell and McDermott 2019). This noise-tolerance allows improved reconstruction of speech sounds and depends on adaptive mechanisms linked to synaptic depression (Rabinowitz, Willmore et al. 2013, Mesgarani, David et al. 2014). However, the majority of studies of sound processing in noise have been conducted in anesthetized or passive subjects, where it is difficult to relate neural function to listening ability.

Understanding auditory cortical processing during behavior is vital for providing context to physiological studies and translating insights from neural circuits to sensory perception. Although many studies report invariance to noise, the improvements in stimulus representations observed in the brain are relatively small compared to large impairments in neural encoding of sounds with decreasing signal-to-noise ratio (SNR). In adverse listening conditions with low SNRs, both neural representation of target sounds and behavioral performance break down (Narayan, Best et al. 2007, Schneider and Woolley 2013, Teschner, Seybold et al. 2016, Christison-Lagay, Bennur et al. 2017) suggesting that degradation in cortical representations may underlie difficulties hearing in noise.

To address the importance of cortical representations for hearing in noise, it is vital to test the causal contribution of auditory cortex. While neural recordings offer valuable insights into the correlations between brain function and behavior, they cannot by themselves show that this function is required for a particular task. Despite the importance of causal tests for our understanding, the need for cortical processing in discrimination of sounds in noise has remained unaddressed. The one study to test the necessity of auditory cortex in noisy listening using lesions indicated an important role in discrimination of frequency-shifted syllables (Porter, Rosenthal et al. 2011). However functional recovery following lesions complicates interpretation of such results, which would benefit from further investigation using inactivation techniques that reversibly suppress auditory cortex.

Here, we addressed the multiple issues of recording and testing the causal contribution of auditory cortex to hearing in noise using an animal model of speech sound (vowel) discrimination in ferrets. Ferrets were trained to discriminate vowels and were tested with and without additive broadband noise while neural activity was recorded or reversibly suppressed using inactivation by cooling. These approaches demonstrated that noise impaired both behavioral and neural discrimination of vowels and that auditory cortex was necessary for sound discrimination in noise. Models of cortical inactivation based on simultaneous recording and cooling of neurons supplemented this work, enabling us to link neural activity to causality through the integration of signals across populations of neurons. Based on these insights, we explored the negative effects of noise on auditory cortical activity to develop strategies for improving auditory behavior. These strategies leveraged the adaptive properties of the auditory system by exposing subjects to continuous background noise. The effects of continuous noise on neural and behavioral discrimination of vowels was then established, while inactivation experiments tested the role of cortex in adaptation to noise. Finally, the translational benefit of such strategies developed in ferrets was tested on human listeners’ ability to hear in noise.

## Results

### A functional role for Auditory Cortex in vowel discrimination in noise

To examine cortical processing of sounds in noise during behavior, we trained ferrets (n = 7) to discriminate synthetic vowel sounds in a two-choice task (Fig. 1A) and tested their discrimination of sounds in clean conditions and when broadband noise was added (Fig. 1B). Animals initiated sounds at a central response spout and were presented with target sounds comprising of a pair of identical vowel tokens (either /u/ or /ε/). Here, two tokens were used for consistency with previous work (Bizley, Walker et al. 2013, Town, Atilgan et al. 2015) but also allowed us to examine neural adaptation within single trials in later experiments. Vowels were either presented alone in clean conditions, or with additive broadband noise that lasted from at onset to the first vowel token sounds to the end of the second token. Vowels were varied in sound level (50 – 80 dB SPL), while noise was presented at a fixed level (70 dB SPL).

**Fig. 1.**
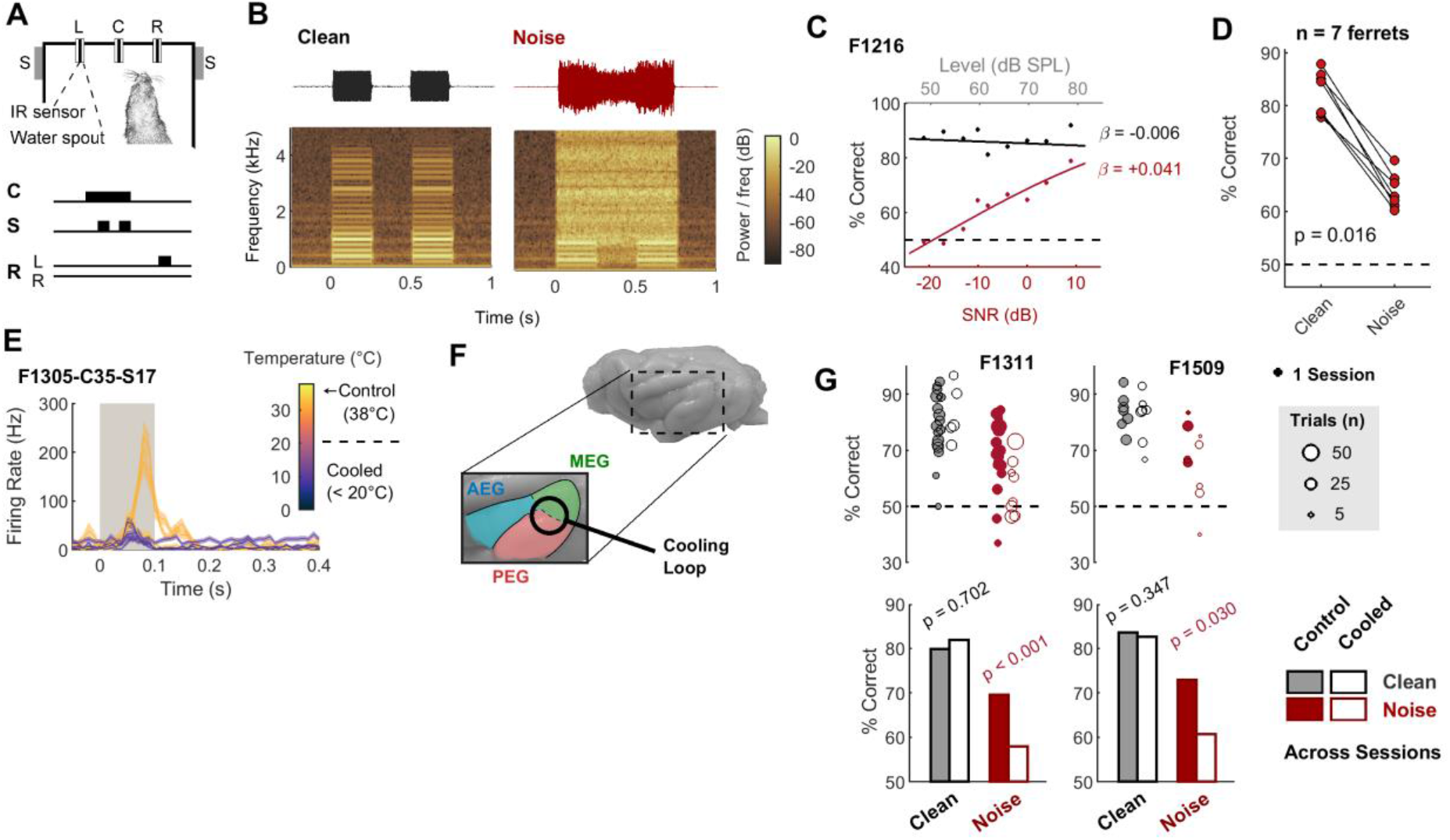
Vowel discrimination in noise. **(A)** Vowel discrimination task and stimulus design. Ferrets initiated trials by visiting a central port (C) and waiting for a variable period before stimulus presentation. Speakers (S) presented sounds (two tokens of the same vowel; blue) to the left and right of the head in all conditions. Animals responded at the left (L) or right (R) port depending on vowel identity. (**B)** Example waveforms and spectrograms recorded in situ showing sounds presented in clean conditions and with additive broadband noise presented in a restricted window beginning at stimulus onset and lasting until the end of sound presentation (0 dB SNR). (**C**) Vowel discrimination performance as a function of sound level (clean) and SNR (noise) for one ferret (F1216). β indicates logistic regression coefficients for models fitted to single trial responses in each condition. (**D**) Performance across SNR or sound level for each ferret (n = 7). Value (p) shows pairwise comparison of performance between clean and noise conditions (sign-rank test). (**E)** Effect of reducing cortical temperature on neural activity. Data shows the mean ± s.e.m. firing rate of one unit recorded from Suprasylvian cortex during presentation of sensory (visual) stimuli in cooled and control conditions. (**F**) Location of cooling loop over boundary between MEG and PEG in auditory cortex for inactivation during vowel discrimination. AEG shows anterior Ectosylvian gyrus region of auditory cortex that was not targeted by cooling loops. (**G)** Performance for each ferret during bilateral cooling or control sessions. Performance shown for each session (top) and across sessions (bottom) with values indicating the probability of observing effects of cooling by chance (permutation test, 10^4^ iterations).

For each animal, we compared vowel discrimination across sound levels (clean conditions) or SNRs (in noise). In clean conditions, task performance was consistent across sound levels (Fig. 1C and Supplementary Figure 1) and significantly better than chance at nearly all signal levels tested for all animals (Binomial test, p ≤ 0.001, Supplementary Table 1). Addition of noise impaired vowel discrimination, with performance across SNR being significantly lower in noise than clean conditions (Fig. 1D, Sign-rank test, median decrease = 18.3%, p = 0.016) and performance at SNRs below −10 dB rarely better than chance (Binomial test, p > 0.05, Supplementary Table 1).

These results suggest that vowel discrimination in noise was dependent on the level of vowel sounds, while performance in clean conditions was largely invariant to sound level. We assessed this by quantifying the effect of sound level on task performance using logistic regression. In noise, all but one ferret (6/7) showed a significant decline in performance with decreased signal level; whereas the same effect was only observed in two animals (2/7) in clean conditions (p < 0.05; Supplementary Table 2). Thus the addition of noise introduced a widespread association between task performance and sound level of target vowels that was rarely visible in clean conditions.

We next asked if auditory cortex was necessary for vowel discrimination by reversibly inactivating this brain region using cooling. Reducing the temperature of cortex suppresses spiking responses of neurons (Fig. 1E), enabling temporary inactivation of the auditory cortex in a single behavioral session (Malhotra, Stecker et al. 2008, Wood, Town et al. 2017). Here, we implanted cooling loops over the low-frequency boundary between middle and posterior Ectosylvian gyrus (MEG and PEG respectively, Fig. 1F) in two ferrets trained to discriminate vowels. Task performance was compared between cooled and control sessions for discrimination of clean or noisy sounds.

Cooling impaired task performance for sounds presented in noise but not in clean conditions (Fig. 1G): For both ferrets tested, performance in noise was significantly worse during cooling than control sessions (F1311: change in performance [control – cooling], ΔC = 11.7%, permutation test, p < 0.001; F1509, ΔC = 12.1%, p = 0.03). Such effects were observed across SNRs (Supplementary Figure 2), while in contrast, there was no significant effect of cooling when discriminating clean sounds for either subject (ΔC < 1%, p > 0.3). Thus auditory cortical function was necessary for normal vowel discrimination only when sounds were presented in noise.

### Cortical encoding of vowels in noise

Our findings indicate that auditory cortex causally contributes towards vowel discrimination in noise, and so we wanted to understand the function of auditory cortical neurons in more detail. We implanted microelectrode arrays in the same region across MEG and PEG as cooling loops and recorded 277 sound responsive units in animals discriminating vowels in clean and noisy conditions (Supplementary Figure 3). We aimed to address three questions: (1) How are auditory cortical responses related to vowel discrimination in quiet and noise? (2) Can the activity of neurons explain the effects of cortical inactivation by cooling on behavior? And (3) can cortical responses to sounds in noise provide insight into how noise impairs vowel discrimination, so that we might develop strategies to improve listening in noise?

To analyse the function of neurons across conditions, we first focussed on 130 units that were responsive to sounds when animals discriminated vowels in both clean and noisy conditions (e.g. Fig. 2A). To find a common metric for the comparison of neural information with animal behavior, we decoded vowel identity from the single trial activity of individual units. Our decoder used a leave-one-out cross validation procedure whereby template responses to each vowel were constructed from responses to all but one trial; the Euclidean distance between this left-out trial and templates was then used to estimate the vowel presented (Town, Wood et al. 2018). The time window of activity considered was variable, with our decoder searching for temporal parameters (start time and duration) for which performance was best. Observed performance was then compared against decoding of shuffled sound identity to highlight units that were significantly informative about vowel identity (permutation test, p < 0.05). With this approach, we found 70/130 units (53.9%) were informative about vowel identity, of which 26/130 units (20.0%) were informative in only clean conditions, 18/130 units (13.8%) in noise only, and 26/130 units (20%) in both conditions (Fig. 2B).

**Fig. 2.**
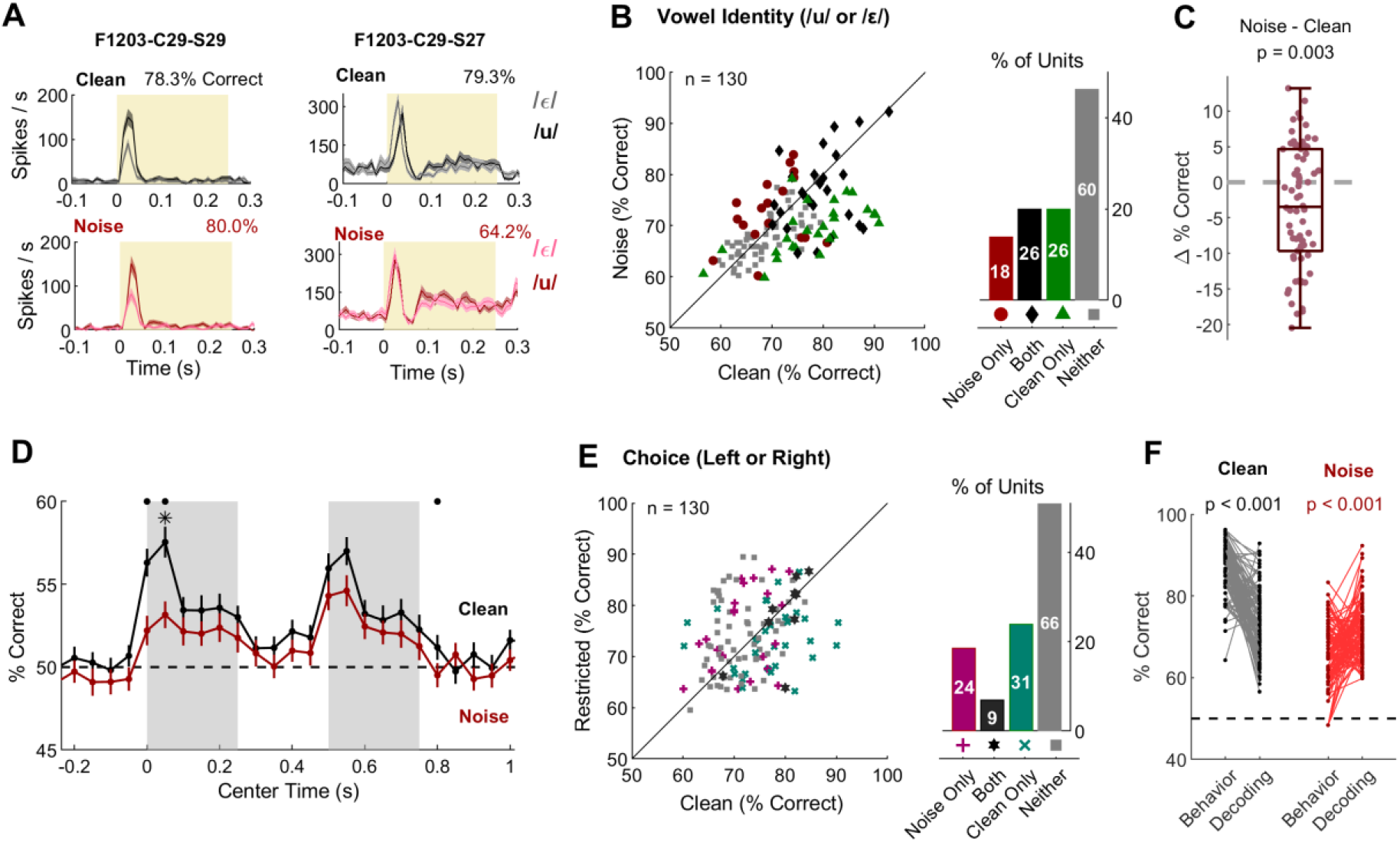
Neural encoding of vowel identity in clean and noise conditions. (**A**) Sound evoked responses of example units to vowels in clean and noisy conditions across sound level and SNR respectively (sound level: 50 to 70 dB SPL, SNR: −10 to 10 dB). Peri-stimulus time histograms show mean ± s.e.m. firing rate. Values indicate optimized decoding performance for sounds in clean and noise conditions. (**B**) Performance decoding vowel identity in noise and clean conditions. Bar plots and markers show units that were significantly informative about vowels in noise (red circles), clean conditions (green triangles), both (black diamonds) or neither condition (grey squares). (**C**) Pairwise difference in decoding performance between clean and noise conditions. Boxplot shows median, interquartile range and 99.3% confidence interval with scatter plot showing individual units. P value indicates effect of condition (sign-rank test). (**D**) Performance decoding vowel identity in a roving 100 ms window: Lines indicate mean ± s.e.m. performance across all units tested (n = 130). Asterisks show significant differences in decoding performance (sign-rank test) with (asterisks) and without (circles) Bonferroni correction for comparison at 31 time points. (**E**) Pairwise comparison of decoding performance with simultaneously observed behavioral performance discriminating vowels in clean and noise conditions. Data shown as in B. (**F**) Performance decoding animal’s choice on single trials in noise and clean conditions. Values (p) indicate significant differences between behavior and decoding (sign-rank test).

As with behavior, decoding performance was worse in noise than clean conditions. When using optimized time windows, decoding performance of units that were informative about vowel identity in either clean or noisy conditions (hereafter “vowel informative units”, n = 70) dropped significantly when noise was added (Fig. 2C; sign-rank test, p = 0.003). Addition of noise also systematically altered the timing optimized decoding windows, typically delaying the best window when decoding sounds in noise (Supplementary Figure 4). These changes in timing parameters suggested that the effects of noise may be temporally specific. This was confirmed using a roving decoding window (Fig. 2D, 100 ms duration) that identified significant noise-related decreases in decoding performance specifically at trial onset (sign-rank test with Bonferroni correction, p < 0.05).

The similarities between effects of noise on behavior and neural decoding prompted us to look more closely at how neural activity related to behavior. We found that it was possible to decode the animal’s behavioral response on individual trials for 65 / 130 units (50%; Fig. 2E, permutation test, p < 0.05), confirming the existence of choice signals in auditory cortex (Niwa, Johnson et al. 2012, Bizley, Walker et al. 2013). However, there were also notable differences in neural decoding and animal behavior. For example, when we compared decoding performance to the behavioral performance of the ferret from the session on which a given unit was recorded, we found significant differences (Fig. 2F): In clean conditions, neural decoding performance was significantly worse than behavioral performance (median difference: −14.9% correct, sign-rank test: p < 0.001); whereas in in noise, behavioral performance was significantly worse than neural decoding (median difference = 5.99% correct, p< 0.001). Similarly, behavioral and decoding performance of individual units was not positively correlated, nor were noise-related changes in behavior and decoding performance (Supplementary Figure 5 and Supplementary Table 3). Thus, while the responses of auditory cortical neurons contained information about both sound identity and choice, such relationships between auditory representation and behavior were only evident at a coarse level.

### Population decoding offers a mechanism for effects of cooling

In primates, population measures of neural function provide a better account of auditory processing in noise than individual units (Christison-Lagay, Bennur et al. 2017). We therefore turned to a population decoding approach to examine the effects of noise on representations of vowel identity. To estimate the identity of vowels from populations of neurons, we summed estimates of vowel identity made in roving windows (0.1 s duration) from 200 units drawn randomly from all 277 sound-responsive units recorded (including but not limited to vowel informative units, as well as units that were only responsive to sounds in noise). Single trial estimates for each unit were made in clean and noise conditions separately, using the same template matching procedure used for individual unit decoding and integrated using a weighted population vector (see Methods). With this approach, we recovered vowel identity in both clean and noise conditions with similar accuracy to the animal’s behavior and with better decoding performance in clean conditions than in noise at trial onset (Fig. 3A).

**Fig. 3.**
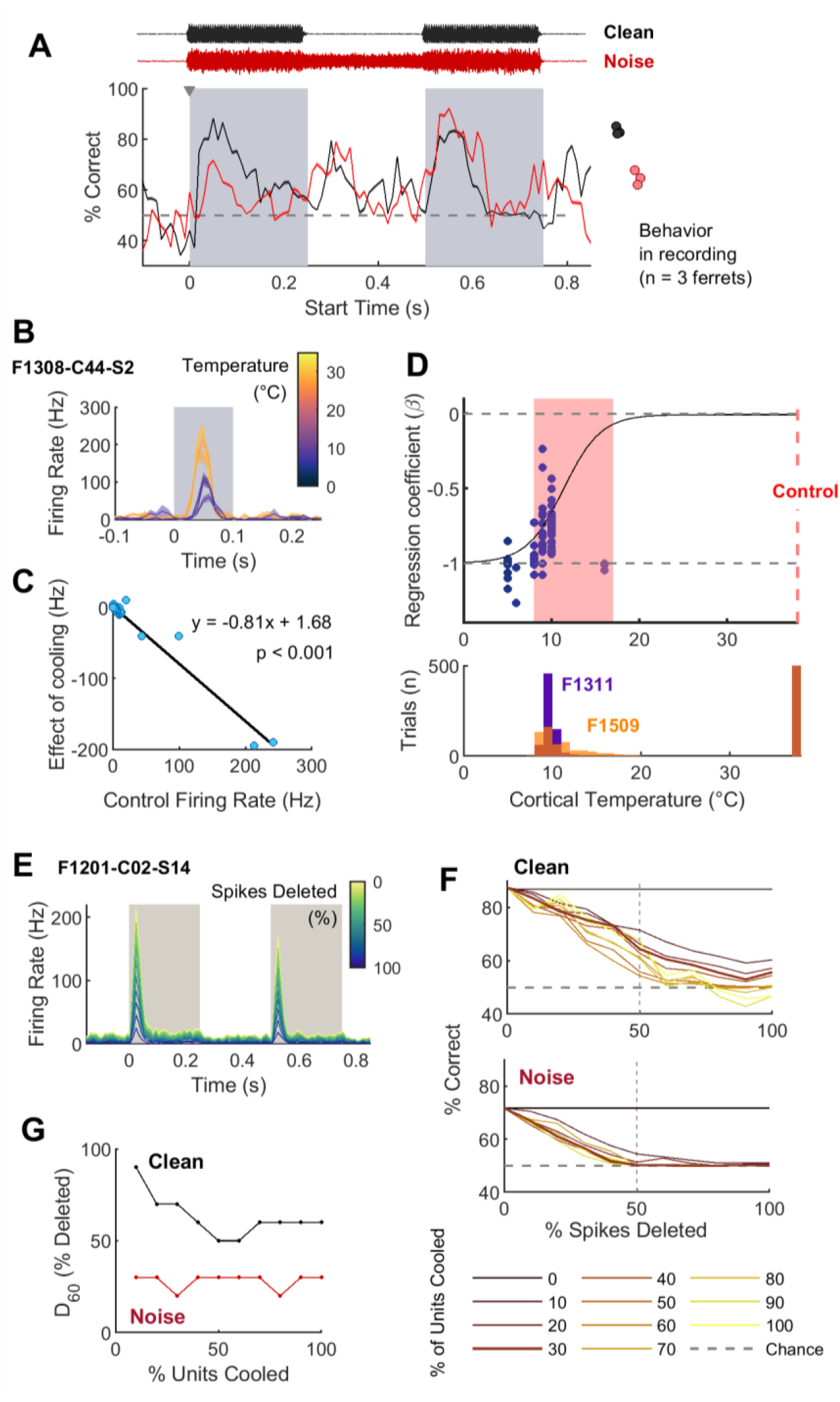
Population decoding and simulations of cortical inactivation. (**A**) Decoding identity of sounds in clean and noise conditions from activity of 200 units using a roving time window (100 ms duration, times plotted with respect to the center of this window). Data shown as mean ± s.e.m. across 100 randomly selected populations comprised of different constituents. Grey triangle indicates time window giving best performance (−50 to 50 ms). (**B**) Responses of one SSY unit to a white light in control (30°C) and cooled (10°C) conditions. (**C**) Comparison of mean firing rate of the same unit as in B during control conditions vs. drop in firing rate with cooling. Dotted line shows linear regression model fitted to firing rates in 10 ms time bins (individual data points). (**D**) Comparison of regression coefficients for 73 units (individual data points) recorded in control and cooled conditions vs. temperature drop achieved during cooling. Line indicates sigmoid estimate the effect of cooling likely to occur during behavior. Red zone indicates range of temperature reductions during behavioral experiments. Histogram shows the distribution of cortical temperatures during cooling recorded in two ferrets during behavior. (**E**) Example response of one unit in auditory cortex during recording (control) and simulated cooling. (**F**) Population decoding performance shown as a function of cooling effect (0 to 100% of spikes deleted) and proportion of units affected (0 – 100% of population) for neural activity sampled between −50 and 50 ms around trial onset. Data shown as mean performance across 100 populations. (**G**) Percentage of spikes deleted to reduce decoding performance below 60% correct (D_60_). Data for F and G shown as mean across 100 simulations.

These results show that population decoding is comparable with behavior when auditory cortex is fully functional, but they also offer new possibilities to understand how cortical inactivation produced selective behavioral deficits in vowel discrimination in noise but not clean conditions. Specifically, because cooling leads to generic reduction in firing rates across cell types (in contrast to pharmacological methods such as muscimol), we could reasonably simulate the effects of cooling on population activity by deleting action potentials from responses of individual neurons. We hypothesized that the reason cooling only impaired vowel discrimination in noise was because information about vowel identity in responses of individual neurons to sounds in noise was scarcer and thus more critical for behavior and population decoding than when clean sounds were presented. In contrast, information (as measured through decoding metrics) about the identity of clean sounds was widespread (and thus redundant) and so behavior and population decoding should be more robust to cortical inactivation.

To simulate the effects of cooling on responses of individual units, we built a simple model mapping suppression of spiking activity to reduction in cortical temperature using data obtained from simultaneous cooling and single unit recording in Suprasylvian cortex (SSY; higher order visual cortex) (Wood, Town et al. 2017, Atilgan, Town et al. 2018). Figure 3B illustrates the suppression of responses of one SSY unit to visual stimuli when cooling was applied. The reduction in spiking activity with cooling was proportional to the original firing rate: peaks in stimulus-evoked responses were most strongly suppressed, whereas baseline activity was less affected (as there were fewer spikes to remove). For each SSY unit, we estimated the dependence between firing rate and effect of cooling using linear regression (Fig. 3C) and compared the gradient of the regression slope with drop in cortical temperature produced by cooling for 73 units (Fig. 3D). We interpreted regression gradients as a proxy for the percentage of spikes suppressed by cooling; values around −1 would reflect complete (100%) inactivation, whereas values near zero would show no effect of cooling. As cortical temperature decreased to lower values, we observed more negative regression slopes that reflected greater reductions in spiking activity. Fitting a sigmoid function to this relationship, we could then work backwards to estimate the proportion of spikes that should be suppressed when cooling auditory cortex during vowel discrimination.

For cortical inactivation during behavior, cortical temperature was reduced from 38°C in control sessions to between 8 and 20°C during cooling (Fig. 3D, mean ± s.d. cortical temperature: F1311: 9.54 ± 1.4°C, F1509: 10.5 ± 2.62°C). This corresponded to a suppression of approximately 50 to 100% of all spikes in the inactivated region of auditory cortex. To implement a simple simulation of cooling, we therefore deleted a proportion of randomly selected spikes from the activity of each unit (e.g. Fig. 3E).

Following spike deletion to simulate effects of cooling, we repeated our population decoding approach either with data from all ‘cooled’ units, or a mixture of cooled and control units that mimicked the anatomical limitations of cooling. Although cooling allows inactivation of large populations of neurons across several millimetres of the cortical surface, ferret auditory cortex encompassed several fields over a larger area (approximately 1 cm^2^). The cooling loops in our study were thus only likely to affect only a third of auditory cortex, in the MEG-PEG border region, and so leave other fields (e.g. AEG) relatively unaffected. The contribution of these unaffected regions may be visible in behavior, where animals’ performance in noise during cooling was impaired, but remained better than chance (Supplementary Table 4). To accommodate these limitations, we retained a sub-population of units without spike deletion that mirrored the contribution of cortical neurons that were not directly affected by cooling. For each set of simulation parameters (proportion of spikes deleted, proportion of units affected) we measured population decoding performance across 100 randomly sampled populations (200 units in each population).

Increasing both the proportion of spikes deleted for each unit and the proportion of units cooled produced larger impairments in decoding performance (Fig. 3F). To draw comparisons with inactivation during behavior, we focussed on decoding performance in the 0.1 s after trial onset, when decoding performance during vowel discrimination was strongest. We found that cortical inactivation was more detrimental to population decoding of sounds in noise than in clean conditions; specifically population decoding of sounds in noise declined to chance levels with fewer units cooled and/or with fewer spikes deleted than decoding sounds in clean conditions. This pattern was reflected in the percentage of spikes we needed to delete in order to reduce population decoding performance below 60% (D_60_) – the level to which behavioral performance declined during cooling. In clean conditions, we needed to delete at least half of all spikes (and often more when inactivating <50% of units) in order to impair performance to this extent, whereas in noise it was possible to do so with inactivation of just 20-30% of spikes. Thus simulated cooling confirmed our hypothesis that neural representation of sound in noise was less robust to cortical inactivation than representations of clean sounds.

### What goes wrong in noise?

The susceptibility of vowel discrimination in noise to auditory cortical inactivation (in vivo or in silico) and its contrast to discrimination in clean conditions emphasizes the salient effect that additive noise exerts on neural processing. This is echoed by our behavioral data showing that at low SNRs, noise substantially disrupts discrimination of vowels that would otherwise be easily differentiated (Fig. 1). Listening in noise represents one of the major challenges for listening in the natural world, and for hearing-impaired patients particularly. We therefore turned to ask if the degradation of auditory representations we observe could show us how noise-related deficits in sound recognition arise.

The primary effect of adding noise to sounds was to increase sound-evoked activity, as one would expect from the addition of broadband energy. Neurons that were already responsive to vowels (n = 130) were driven even more strongly, with higher average firing rates in clean than noise conditions (Fig. 4A). Noise-induced increases in firing rates were particularly evident at trial onset, where a significant increase was observed across units at onset of the first vowel token (Fig. 4B; sign-rank test with Bonferroni correction for comparison at multiple time points, p < 0.005). In contrast, neurons became significantly less responsive to the second vowel token, indicating a potential adaptation to noise across the trial (p < 0.005). This is consistent with decoding performance, where recovery of vowel identity in noise occurred later than in clean conditions (Supplementary Figure 4) and imply that longer exposure to noise may have improved decoding and behavioral performance.

**Fig. 4.**
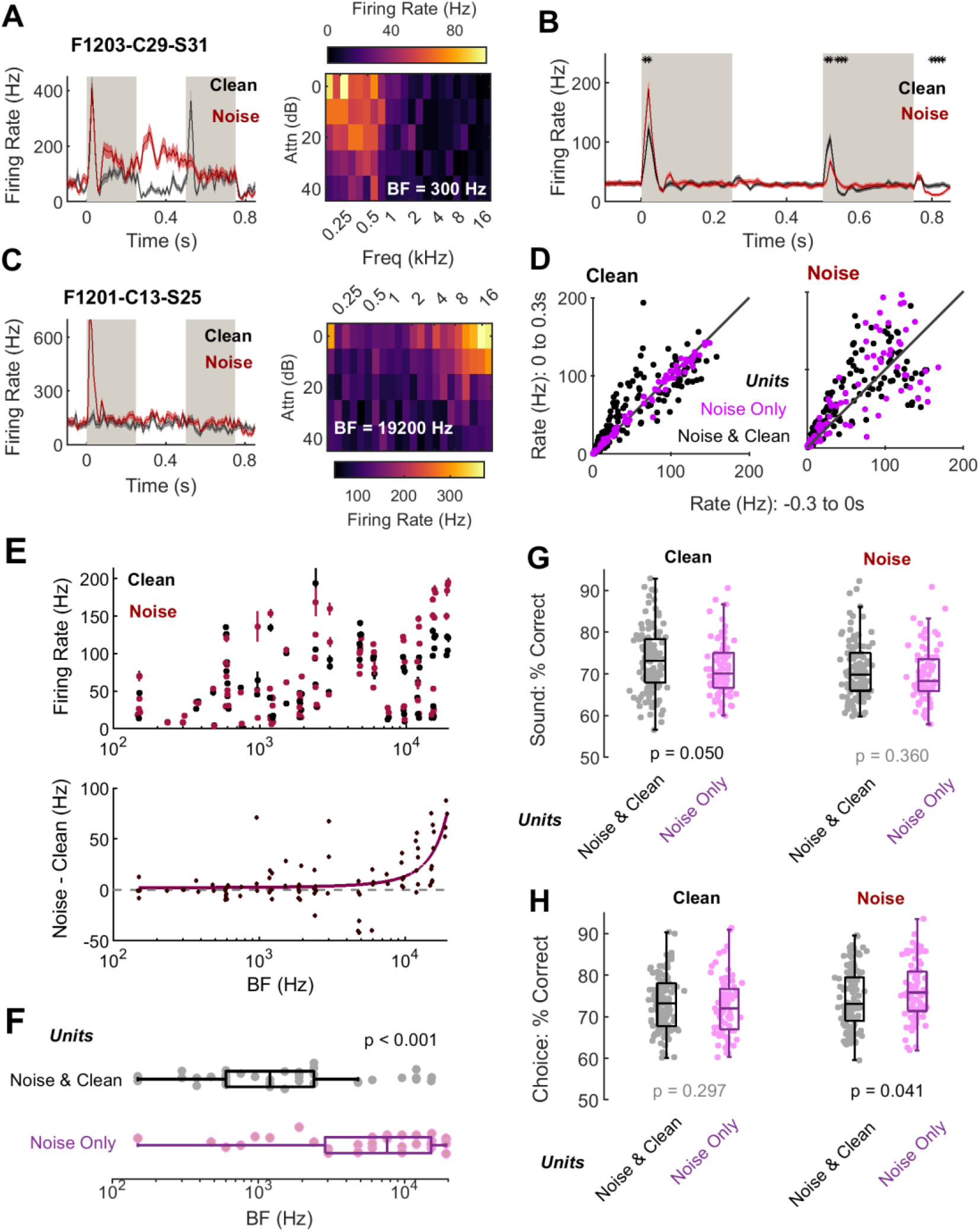
Noise-related changes in cortical responses to vowels. (**A**) Example responses to vowels and frequency tuning recorded from an example units that responded to vowels in both clean and noise conditions. BF indicates best frequency of units. Firing rate shown as mean ± s.e.m. across trials. (**B**) Mean ± s.e.m. firing rate across all units (n = 130) that were responsive to vowels in clean and noise conditions. Data shown as mean ± s.e.m; markers indicate significant differences between conditions (sign-rank with Bonferoni correction, p < 0.05). (**C**) Example response and frequency tuning of a unit that was only responsive to sounds in noise. Data shown as in A. (**D**) Pairwise comparison of firing rates for each unit used to assess responsiveness to vowels in clean and noise conditions. Data points show mean firing rate for each unit in the before (−0.3 to 0 s) or after (0 to 0.3 s) trial onset. Data points are coloured according to whether the unit responded in both clean and noise conditions, or only in the noise condition. (**E**) Mean sound-evoked response to vowels in clean and noise condition plotted vs. best frequency for 89 units in which responses to tones and vowels were both measured. Top: Data for responses to sounds in noise and clean conditions separated with error bars showing standard error across trials. Bottom: Pairwise comparison with exponential model fitted to effect of noise (purple line), indicating that higher frequency units were more sensitive to broadband noise. (**F**) Comparison of frequency tuning of units that were responsive to vowels in both clean and noise conditions, or only for vowels in noise. Box plots show median and interquartile range of BFs for units that were responsive to vowels only in noise (red, n = 33) or in both conditions (black, n = 44). Value (p) indicates unpaired comparison of CF across groups (rank-sum test, p = 2.24 x 10-5). (**G-H**) Performance decoding vowel identity (G) or behavioral choice (H) in clean and noise conditions, shown separately for units that responsive only in noise, or in both conditions. Box plots show median and interquartile range with individual data points showing each unit. Values (p) show rank-sum comparisons across sub-groups of units (noise only vs. noise & clean).

A larger population of neurons was also active when sounds were presented in noise: In 30% of neurons (84 / 277 units), vowels in noise elicited significant responses that were not present in clean conditions (Fig. 4C-D). Such units were tuned to higher sound frequencies than units driven by vowels across conditions (Fig. 4E-F; rank-sum test, p < 0.001), indicating that the inclusion of broadband energy in the noise may have recruited a population of cells with spectral tuning that overlapped less with the spectra of the speech sounds (which contain most energy at frequencies below 4 kHz). Dividing units by their responsiveness also revealed functional differences: In clean conditions, units that were only responsive to sounds in noise showed significantly poorer decoding of vowel identity compared to units that were responsive across conditions (Fig. 4G; rank-sum, p = 0.05). In contrast, there was no difference between these groups of units when decoding the identity of sounds in noise (p = 0.360).

Interestingly, when noise was added to sounds, choice-probabilities (as measured by ability to decode animal’s behavioral responses) were significantly higher in the noise-only sub-group of units than in units that responded across conditions (Fig. 4H, rank-sum test, p = 0.041). Choice decoding for noise-only units was also significantly higher for responses to sounds in noise than in clean conditions (sign-rank, p < 0.001). This suggests that not only did the addition of noise to the stimulus drive these neurons, but their activity became more closely linked to the animal’s behavior. In contrast, the choice probabilities for units that were responsive across conditions did not change when noise was added to sounds (sign-rank, p = 0.300).

Units that only responded to sounds in noise could provide an independent estimate of the properties of background noise, and thus offer a signal for denoising representations of target sounds. We did find examples of noise-tolerant representations of sound identity within auditory cortex (Supplementary Figure 6), although the information about vowel identity preserved in the responses of these units would not explain why noise impairs vowel discrimination during behavior. Instead, our results suggest that activity of noise-only units was integrated into decision making when discriminating vowels in noise, with potentially negative effects on task performance.

Our results suggest that addition of noise to sounds produces excessive drive of neurons at trial onset, which coupled with recruitment of otherwise unresponsive neurons, may impair vowel discrimination. If true, task performance may be recovered if the excitatory effects of noise at trial onset were reduced, and noise-only units were inactivated. Cortical inactivation using methods such as cooling cannot target functionally defined neural populations, and while sophisticated optogenetic techniques exist to do so (Mardinly, Oldenburg et al. 2018), such methods are currently limited in clinical applicability for widespread use in humans. We therefore turned to a simpler approach to inactivate neurons, using the acoustic properties of sounds, combined with the adaptive properties of the auditory system (Perez-Gonzalez and Malmierca 2014, Willmore, Cooke et al. 2014) to suppress noise-related activity by pre-exposure to noise.

### Continuous exposure improves sound discrimination in noise

Adding noise to sounds impaired vowel discrimination behavior while also driving additional spiking activity in auditory cortex, both in neurons that already responded to vowels and newly recruited units. The effects of noise diminished across the trial: unit responses to noise were often limited to trial onset, and the effects of noise on population spiking and decoding performance declined with time - being smaller or absent at second vowel token compared to first token (Fig. 4A). These results suggest that cortical activity may rapidly adapt to sounds and that behavioral impairments in vowel discrimination may be linked to changes in coding at sound onset. One way to potentially counter impairments to task performance in noise may thus be to pre-expose subjects to noise and so restore normal responses to vowels at trial onset by providing listeners with opportunity to adapt to listening conditions before sound presentation. We therefore contrasted auditory cortical responses to sounds during vowel discrimination in clean and noise conditions with responses to sounds discriminated in continuous noise.

Continuous noise (70 dB SPL) was presented to the same animals implanted with electrodes in auditory cortex while they discriminated vowel sounds. Continuous noise shared the same broadband spectrum as temporally restricted noise that was only present with vowel sounds (Fig. 5A), but was presented from the beginning to the end of each behavioral session (10 – 45 minutes). We recorded 128 units in continuous, clean and temporally restricted conditions (Fig. 5B), of which 41% (53 / 128) were responsive to vowels in all conditions. Of the remaining units that were only responsive in specific conditions, significantly fewer were responsive in continuous than restricted noise (29 vs 43 units, Χ^2^ = 5.24, n = 75, p = 0.022), indicating that continuous exposure successfully countered noise-related recruitment of additional responsive units. This was also evidenced by population firing rate of units recorded across conditions, where continuous exposure to noise abolished the enhanced response at trial onset observed when restricted noise was added (Fig. 5C).

**Fig. 5.**
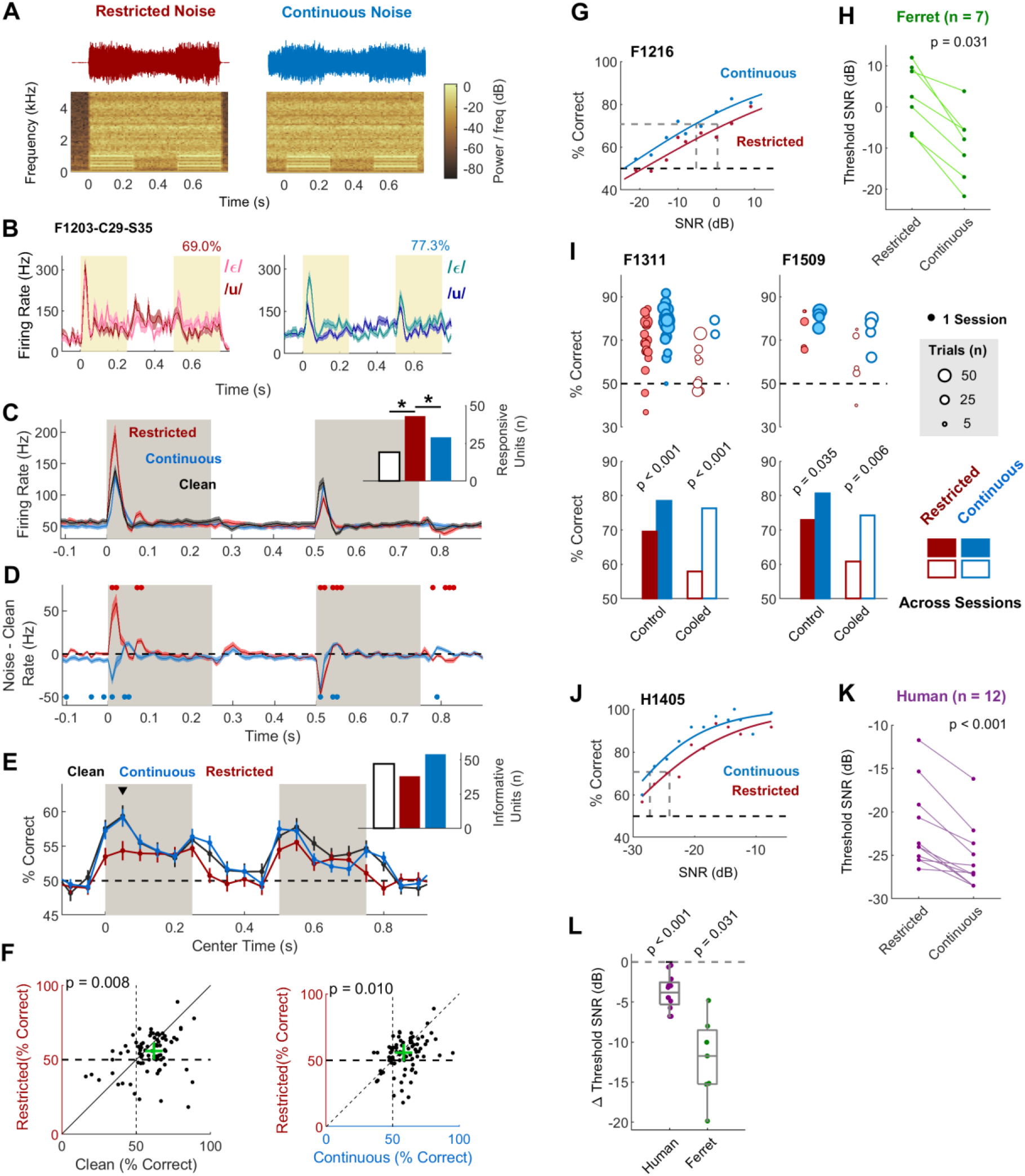
Continuous exposure to noise reshapes cortical processing and improves sound discrimination. **(A**) Stimulus waveforms and spectrograms illustrating vowels in continuous and temporally restricted noise (0 dB SNR). (**B**) Sound evoked responses from one example unit to vowels in continuous and restricted noise (across SNR: −10 to 10 dB). Data shown as mean ± s.e.m. firing rate. Percentage values indicate performance decoding vowel identity. (**C**) Mean firing rate across 128 sound-responsive units recorded in clean, continuous and restricted noise conditions. Data shown as mean ± s.e.m across units. Inset shows the proportion of units that were responsive to sounds by condition. Asterisks indicate significant differences in number of responsive units (Chi-squared test, p < 0.05). (**D**) Change in firing rate from clean to restricted or continuous noise across all units (n = 128). Data show mean ± s.e.m. difference in firing rate (clean – noise condition). Markers indicate times at which effects of noise differed significantly from zero (sign-rank test, Bonferroni corrected for 100 comparisons, p < 0.005). (**E**) Performance decoding vowel identity in a roving 100 ms window: Lines indicate mean ± s.e.m. performance across all units that were informative about vowel identity in clean or either noise conditions (n = 83). Inset shows the number of units that were significantly informative about vowel identity in each condition. Black triangle indicates time of comparison shown in scatter plots below in F. (**F**) Pairwise comparison of decoding performance in clean conditions vs. restricted noise (left), and continuous noise vs. and restricted noise (right). P values indicate sign-rank comparisons; green cross-hairs show median across all informative units. (**G**) Performance of one ferret (F1216) discriminating vowels at each SNR (individual data points) in continuous and restricted noise. Full lines show logistic regression models fitted separately for each noise type; dashed lines indicate performance at which thresholds were calculated (grey) and chance performance (black). (**H**) Thresholds measured at 70.7% correct for all ferrets (n = 7) with sign-rank comparison in continuous vs. restricted noise (p). (**I**) Performance for two ferrets during bilateral cooling and control sessions. Performance shown for each session (top) and across sessions (bottom) with values (p) indicating the probability of observing effects of cooling by chance (permutation test, 10^4^ iterations). (**J**) Performance of one human (H1405) discriminating vowels at each SNR in continuous and restricted noise. Data shown as in G. (**K**) Thresholds measured at 70.7% correct for all humans (n = 12) with sign-rank comparison in continuous vs. restricted noise. (**L**) Change in threshold (Continuous – Restricted) for humans and ferrets. Individual data points show each subject; box plots show median and inter-quartile range. Values (p) indicate significant change in threshold vs. zero (sign-rank test).

To further contrast the effects of noise type on responses to target vowel sounds, we subtracted the response of each unit to vowels in clean conditions from responses to sounds in continuous and restricted noise (Fig. 5D). This revealed that auditory responses in continuous noise were significantly weaker than to vowels in even clean conditions at trial onset (sign-rank test with Bonferroni correction, p < 0.005). We also observed significant differences in pre-stimulus activity that suggest continuous exposure to noise produced small but long-lasting suppression of cortical activity. The effects of continuous noise at trial onset were contrasted starkly with the effects of restricted noise, which significantly enhanced responses to onset of the first vowel token. However, by the onset of the second vowel token, vowels in both continuous and restricted noise generated significantly weaker responses than in clean conditions. This suggests that adaptive processes engaged by exposure to noise may be rapid and operate at the timescale of hundreds of milliseconds, such that exposure to 500 ms of noise is equivalent to exposure for several seconds or minutes.

Continuous exposure to noise also improved our ability to decode vowel identity from neuronal response to sounds in noise (Fig. 5E): More units were significantly informative about identity in continuous (54/128 units; 42.1%) than temporally restricted noise (38/128 units; 29.7%), while decoding of vowel in continuous noise occurred earlier and over longer time windows (Supplementary Fig. 6) indicating that information was available for longer in the trial. The longer availability of information in continuous noise was echoed by comparison of decoding performance across trial, where exposure to continuous noise resulted in improvements in decoding relative to restricted noise at nearly all time points (Fig. 5E). This improvement was significant when considering the time window at trial onset (n = 83 vowel informative units, median improvement = 2.94%, sign-rank test, p = 0.010), when the addition of restricted noise had produced significant impairments in decoding performance when compared to clean conditions (median impairment: 4.56%, p = 0.008). In contrast, there was no significant difference in decoding performance in clean or continuous noise conditions (p > 0.05). Thus continuous exposure to noise restored the availability of information about sound identity in cortical activity, including at the potentially critical time at trial onset.

Improvements in performance decoding neural activity were mirrored by behavioral benefits of continuous exposure to noise. Vowel discrimination in clean and restricted noise was tested for the seven ferrets that also performed the task in clean conditions. For all subjects, we observed better task performance in continuously than restricted noise (Fig. 5G and Fig. S1). At a group level, this was reflected by a significant reduction in thresholds in continuous compared to restricted noise (Fig. 5H, 70.7% correct thresholds, Wilcoxon sign-rank test, n = 7, *p* = 0.031).

To understand if changes in cortical processing to sounds in continuous noise were the driving force for improvements in behavior, we conducted further inactivation experiments by cooling auditory cortex in two ferrets discriminating sounds in restricted and continuous noise. In both control and cooled sessions, performance was significantly better in continuous than restricted noise in each subject tested (Fig. 5I; F1311: change in performance [continuous – restricted], ΔC = 18.4%, permutation test, p < 0.001; F1509: ΔC = 13.5%, p = 0.006) indicating that the effects of continuous exposure on behavior were not cortically-dependent. Indeed the performance benefit of continuous noise exposure was larger during cooling than in control sessions (change in performance: [continuous _Cooled_ – restricted _Cooled_] - [continuous _Control_ – restricted _Control_], F1311: +8.85%, p = 0.0002; F1509: ΔC = +7.69%, p = 0.035). This effect was driven by the poor performance of animals when discriminating sounds in noise during cooling: performance in restricted noise was worse in cooling than control sessions (Cooled – control [ΔC], F1311: ΔC = −11.7%, permutation test, p < 0.001; F1509, ΔC = −12.1%, p = 0.03), whereas only in one animal was performance in continuous noise impaired by cooling (F1509, ΔC = −6.35%, p = 0.022; F1311: ΔC = −2.12%, p = 0.266). Thus, while we identified a causal contribution of auditory cortex to sound discrimination in noise, cortical inactivation did not suppress the benefits of continuous noise exposure on task performance.

Finally, we asked if the behavioral benefits of exposure to continuous noise observed in ferrets translated to human listeners. Human subjects (n = 12; 7 male) were tested in a vowel discrimination task designed to match the task performed by ferrets as closely as possible. Vowels were presented in the same temporally restricted or continuous noise, although not in clean conditions during testing. Like ferrets, humans performed better in continuous than restricted noise (Fig. 5J and Supplementary Fig. 1) and continuous exposure to noise significantly reduced performance thresholds (Fig. 5H, thresholds calculated at 70.7%, sign-rank test, *p* = 4.88 x 10^-4^). Thus vowel discrimination by both humans and ferrets benefited from continuous exposure to noise, in conditions in which we also observed improvements in auditory cortical processing of vowels in continuous noise in ferrets.

## Discussion

We found that the addition of broadband noise to sounds impaired the ability of ferrets to discriminate vowel identity and that auditory cortex makes a causal contribution to sound discrimination in noise. Noise-related impairments in behavior were mirrored by reduction in performance decoding vowel identity from responses of auditory cortical neurons, both at the level of individual units and neural populations. Modelling and population decoding showed how this reduction in information could produce behavior that was more susceptible to cortical inactivation. The addition of noise altered cortical processing by increasing firing rates in vowel-responsive units and recruiting additional neurons that were not responsive to vowels alone. The temporal profile of these effects suggested that adaptation may be used to counter the effects of noise on cortical processing and behavior. We found evidence supporting this as exposing subjects to continuous background noise suppressed the effects of noise on cortical activity while also improving decoding and behavioral performance. The behavioral benefits of adaptation to noise were observed in both ferrets and humans but were not dependent on auditory cortical function.

### Cortical inactivation

Our study shows an important functional contribution of auditory cortex to speech sound discrimination in noise. While inactivation studies have consistently shown a role for auditory cortex in sound localization (Smith, Parsons et al. 2004, Malhotra, Stecker et al. 2008), support for involvement in other auditory behaviors remains equivocal (Talwar, Musial et al. 2001, Cooke, Zhang et al. 2007, Aizenberg, Mwilambwe-Tshilobo et al. 2015, Gimenez, Lorenc et al. 2015, O’Sullivan, Weible et al. 2019). Causal tests of auditory cortex in behavior are critical because studies of auditory cortical function during sound discrimination have been mostly limited to correlational analyses (Bizley, Walker et al. 2013, Mesgarani, David et al. 2014, Engineer, Rahebi et al. 2015, Town, Wood et al. 2018). In one of the few examples to address the causal role of auditory cortex to sound discrimination in animals, Porter et al. (2011) found that following initial deficits in performance following auditory cortical lesion, rats could discriminate frequency-shifted syllables in both clean and noise conditions. This contrasts with our finding of selective deficits in vowel discrimination in noise but not clean conditions. The differences in results may be partly explained by post-lesion recovery of function that is avoided when using techniques to reversibly inactivate neural tissue; however there are also important differences in stimuli, task design and model species that further complicate comparison across studies.

When discriminating sounds in clean conditions, coding of vowel identity across the population was highly redundant and thus robust to partial inactivation by cooling. Our simulations predict that cortical inactivation would be effective only when the spiking activity of most / all neurons is completely disabled. Although it would be ideal to control inactivation across the entire area of auditory cortex, the size and geometry of this brain area in ferrets (which surrounds and extends into the ectosylvian sulcus) makes complete inactivation technically challenging. Lesions offer one way to permanently remove large cortical areas, but are irreversible and susceptible to functional recovery. Pharmacological techniques (e.g. application of muscimol) offer a reversible alternative but, like cooling, are unlikely to completely inactivate the large cortical volume that auditory cortex covers. Advanced optical approaches may enable larger coverage and temporally specific inactivation of brain areas (Guo, Li et al. 2014), however additional steps are needed to apply such techniques to freely-moving ferrets, and even then, may not inactivate a sufficient proportion of cells within redundant networks.

In the comparison of vowel discrimination in clean and restricted noise conditions, it may be more accurate to consider clean sounds as a favourable case of continuous noise with high SNR, where noise is not presented to subjects explicitly but rather exists within the environment at some very low level. From this perspective, the effects of cooling we observe on vowel discrimination in noise reflect the contribution of auditory cortex in low SNR conditions. If true, future experiments that reduce the sound level of clean vowels may also reveal causal contributions of auditory cortex.

### Signal processing in noise

Our study used broadband noise, whose statistics remained stationary across trials and sessions; however the selection of noise properties can have important implications for studies of auditory processing. In ferret auditory cortex, reconstruction of speech sounds from neural activity tend to be better in pink (1/f) than white (broadband) noise (Mesgarani, David et al. 2014). This suggests that energy at high sound frequencies may be an important negative influence on neural responses, and is consistent with our findings that the addition of broadband noise recruited units tuned to higher frequencies that were linked to animals’ behavioral choices. It is interesting that the engagement of these units, that were only responsive in noise, was associated with deficits rather than improvements in vowel discrimination. Such results indicate that downstream neurons integrating signals from auditory cortex sample responses from a wide population of cortical neurons rather than a select group of cells that were responsive to target sounds across clean and noise conditions. From a noise-cancelling perspective, the activity of units that are only responsive to noise offers a channel through which to independently estimate and subsequently remove background noise. Here, we initially trained animals to discriminate sounds in clean conditions; however it may be possible to instigate such noise cancellation strategies by training animals to discriminate sounds in noise.

Several studies have reported noise-invariant representations of sound in auditory cortex that could support robust recognition of sounds in noise (Ding and Simon 2013, Rabinowitz, Willmore et al. 2013, Mesgarani, David et al. 2014, Kell and McDermott 2019). However, behavioral results from our study and others (Hienz, Aleszczyk et al. 1996, Shetake, Wolf et al. 2011, Schneider and Woolley 2013) show that recognition of sounds in noise is only robust at high SNRs and breaks down at low SNRs. This suggests that noise-invariant representations may not always give rise to invariant behavior, or fail as SNR decreases in increasingly adverse listening situations. Indeed, when zebra finches fail to discriminate songs against a background chorus at low SNRs, neural responses appear more like the chorus than song (Schneider and Woolley 2013). Although we could identify noise-tolerant representations in ferret auditory cortex (Supplementary Figure 6), the response properties of these cells does not account for the breakdown in behavioral performance that we observed simultaneously with neural recordings.

### Origins and consequences of adaptation to noise

Our findings are consistent with recordings in human auditory cortex (Khalighinejad, Herrero et al. 2019) where ongoing exposure to noise leads to suppression of cortical activity and enhanced neural and perceptual representation of speech sounds. However, our cortical inactivation results suggest that adaptation observed within cortex may not be cortical in origin, but rather may arise earlier in the ascending pathway. Adaptation has been noted in multiple subcortical structures including auditory nerve fibres (Costalupes, Young et al. 1984, Perez-Gonzalez and Malmierca 2014, Willmore, Schoppe et al. 2016) and may even reflect an inheritance of the biophysical properties of hair cell mechanotransduction (Kennedy, Evans et al. 2003). In future, it may be interesting to target the specific cellular mechanisms responsible for adaptation in earlier areas of the auditory system to identify the locations responsible for the behavioral improvement in sound discrimination we observe.

The time course of adaptation to noise appeared to be rapid, on the sub-second level. Although our use of continuous and temporally restricted noise did not provide a detailed temporal comparison, we noted that neural responses to the second vowel token were significantly weaker in restricted noise than clean conditions. This indicated that adaptation had occurred across the trial, within 0.5 – 0.6 seconds of noise onset (or possibly sooner since this measure includes a 0.25 s interval between vowel tokens). This time-scale is consistent with earlier examples of adaptive processes within auditory cortex (Phillips 1985, Ulanovsky, Las et al. 2004).

Adaptation in neural activity across the trial to temporally restricted noise appeared to provide little behavioral benefit, even though adaptation was also associated with improved behavioral performance in continuous noise. The reason for this is likely due to the time-course of decision making in our behavioral paradigm; perception of vowel timbre is associated with acoustic processing at sound onset (Walker, Bizley et al. 2011, Town, Wood et al. 2018) and animals in our task were only required to remain at the central spout during the first vowel token. It is likely therefore that animals weighted their decisions strongly towards acoustic signals at trial onset, when adaptation within the trial had yet to occur and thus temporally restricted noise was most disruptive. Nonetheless, the recovery in behavioral performance and normalization of spiking activity with continuous noise exposure suggests that hearing in noise does benefit from adaptation within the auditory system, so long as listeners have sufficient exposure to noise signals. Additional experiments in which restricted noise is introduced with a range of temporal onsets relative to the vowel sounds will be required to characterize the precise time-course of adaptation and its potential benefits for sound discrimination.

### Relevance for hearing loss

The challenges of hearing in noise are particularly problematic for individuals with hearing loss or deafness and those using hearing aids or cochlea implants, who receive limited benefits from noise-reduction techniques (Brons, Houben et al. 2012, Chong and Jenstad 2018). Approaches to reduce noise often attempt to characterize and subtract background sounds from incoming signals, essentially mimicking the effects of adaptation observed in neural responses to ongoing sounds. However, our findings imply that filtering out background noise may be harmful for performance of hearing aids and cochlea implants, particularly if this deprives neurons in the central auditory system of the chance to adapt to noise, or introduces common onsets for speech sounds and noise used in our study. Although counter-intuitive, it may thus be valuable to intentionally expose listeners to (at least low levels of) background noise in order to prospectively engage adaptive mechanisms and improve speech recognition. Similar suggestions have also been made for leveraging benefits of stochastic resonance using noisy stimuli (Morse and Evans 1996) and so future experiments must assess the audiological value of physiologically inspired strategies for restoring hearing in noise.

## Methods

### Subjects

In behavioral experiments, we trained and tested seven ferrets (0.5-5 years old) in a vowel discrimination task. Subjects were water-restricted prior to testing and received a minimum of 60ml/kg of water either during task performance or supplemented as a wet mash made from water and ground high-protein pellets. Subjects were tested in morning and afternoon sessions on each day for up to five days in a week. The weight and water consumption of all animals was measured throughout the experiment. Regular otoscopic examinations were made to ensure the cleanliness and health of ferrets’ ears. Animals were maintained in groups of two or more ferrets in enriched housing conditions. In anesthetized experiments in which we determined the impact of cortical cooling, we used a further four naïve ferrets (1-2 years old) with no training in any behavioral task. Ferrets used in all experiments were pigmented females. All experimental procedures were approved by local ethical review committees (Animal Welfare and Ethical Review Board) at University College London and The Royal Veterinary College, University of London and performed under license from the UK Home Office (Project License 70/7267) and in accordance with the Animals (Scientific Procedures) Act 1986.

We also tested 12 human subjects (7 male; mean ± s.d. age = 23 ± 1.96) that had no previous experience of the specific vowel sounds used in the study. No subjects reported hearing loss or impairment. All subjects provided written informed consent in accordance with the Declaration of Helsinki. Ethical approval was provided by the UCL Research Ethics Committee.

### Stimuli

Vowels were synthesized in MATLAB (MathWorks, USA) using an algorithm adapted from Malcolm Slaney’s Auditory Toolbox (https://engineering.purdue.edu/~malcolm/interval/1998-010/ that simulates vowels by passing a click train through a biquad filter with appropriate numerators such that formants are introduced in parallel. In the current study, four formants (F1-4) were modelled: /u/ (F1-4: 460, 1105, 2857, 4205 Hz), /ε/ (730, 2058, 2857, 4205 Hz), /a/ (936, 1551, 2975, 4263 Hz) and /i/ (437, 2761, 2975, 4263 Hz). For behavioral experiments (including cortical inactivation and neural recording), ferrets and humans were only trained to discriminate two vowels: either /e/ and /u/ (five ferrets, seven humans), or /a/ and /i/ (two ferrets, five humans). All vowels were generated with a 200 Hz fundamental frequency and roved in sound level to vary SNR.

Vowels were presented in clean conditions as two tokens (250 ms duration) of the same identity, separated with an interval of 250 ms. Vowels were also presented with additive broadband noise fixed at 70 dB SPL. Noise was initially restricted to the time window of stimulus presentation (0 to 750 ms after onset of the first vowel token) and generated afresh on each trial. Onsets of both vowel sounds and noise were independently ramped using a 5 ms cosine function. We also presented vowels in continuous noise that lasted the entire length of the testing session (10 – 40 minutes).

Sound stimuli were presented through two loudspeakers (Visaton FRS 8) positioned on the left and right sides of the head at equal distance and approximate head height. These speakers produce a smooth response (±2 dB) from 200Hz to 20 kHz, with a 20 dB drop-off from 200 to 20 Hz when measured in an anechoic environment using a microphone positioned at a height and distance equivalent to that of the ferrets in the testing chamber. All vowel sounds were passed through an inverse filter generated from calibration of speakers to golay codes.

We also measured the frequency tuning of neurons using sinusoidal pure tones that ranged in frequency from 150 Hz to 19200 Hz in 1/3 octave intervals and sound level from 0 to 30 or 40 dB attenuation. Each tone was presented for 100 ms with a 5 ms cosine ramp and random interval (minimum 250 ms) between tones. Sound levels for all pure tone sounds used to measure frequency tuning were calibrated independently to ensure a fixed sound level for a given attenuation. Stimulus generation and presentation in behavioral tasks were automated using custom software (Matlab, R2013b) running on personal computers, which communicated with TDT real-time signal processors (RM1; Tucker-Davis Technologies, Alachua, FL) and animal (RZ6; TDT) experiments.

### Task design for Ferrets

Ferrets were trained to discriminate the synthetic vowel sounds within a custom-built double-walled sound attenuating chamber (IAC) lined with acoustic foam. The chamber contained of a wire-frame pet-cage with three ports containing infra-red sensors that detected the ferret’s presence (Fig. 1A). On each trial the animal was required to approach the center spout and hold head position for a variable period (0 – 500 ms) before stimulus presentation. Animals were required to maintain contact with the center spout until 250 ms after the presentation of the first token, at which point they could respond at left or right response ports. Correct responses were rewarded with water while incorrect responses led to a brief time out (3 - 8 s).

Ferrets were initially trained to discriminate vowels that roved in sound level over a 6 to 12 dB range (Town, Atilgan et al. 2015). Sounds were subsequently varied over larger ranges of sound levels, as well as fundamental frequency, location and voicing (Town, Wood et al. 2018). Animals were considered trained once they could consistently discriminate vowels with ≥ 70% accuracy across multiple days. Ferrets were then tested by presenting vowels with varying sound level in clean conditions, or with the addition of restricted or continuous noise. Animals were tested on clean sounds and sounds in temporally restricted noise on different trials within the same behavioral session; however testing in continuous noise was conducted in separate sessions usually on the same or subsequent day.

### Task design for Humans

Training and testing was designed to match, as closely as possible, the task design used for ferrets: Subjects performed a two-choice task, seated in a sound isolating booth, in front of a laptop with two speakers positioned at approximately 45° to the left and right of the head. On each trial, the participant initiated sound presentation by key press and was presented with two tokens of a vowel sound, each lasting 250 ms, with a 250 ms interval. The identity of the vowel was constant within trial, and subjects were required to respond using left/right arrow keys in order to receive feedback on whether the response was correct or incorrect. Incorrect responses led to correction trials in which the previous trial was repeated until a correct response was given.

To minimize differences between human and non-human experiments, subjects were given no explicit instructions beyond details of the trial structure. Like ferrets, human participants therefore had to initially learn, by trial and error, which vowel was associated with which response. Training consisted of blocks of 30 (non-correction) trials in which each vowel was presented at one of three fundamental frequencies (200, 330 or 499 Hz), with a fixed sound level (60 dB SPL). Performance was measured as the percentage of correct responses (excluding correction trials) and was assessed at the end of each block. Participants graduated from the training stage when block performance exceeded 85% correct.

After training, subjects completed 12 test blocks in which vowels were presented in continuous or temporally restricted noise. Each vowel was presented at one of six fundamental frequencies (149, 200, 23, 330, 409 or 499 Hz) while sound level was varied between 42 and 62 dB SPL (SNR = −28 to −8 dB). Correction trials were not used during the test phase of the study.

### Behavioral data analysis

To generate psychophysical curves for humans and ferrets, we fitted a logistic regression model that used SNR to predict the proportion of trials performed correctly. Only SNR values for which subjects made 20 or more responses were included in the analysis. Models were fit separately to subjects’ responses to sounds in restricted and continuous noise (and for ferrets, also with sounds presented in clean conditions). Regression models were then used to estimate the association between SNR and performance, as well as threshold SNR values that would produce performance of 70.7%. Thresholds for all subjects (ferrets or humans) were compared across conditions (e.g. restricted vs. continuous noise) using a sign-rank test.

### Reversible inactivation by cooling

Cortical inactivation experiments were performed using an approach developed by Wood et al. (2017): Two ferrets were implanted with cooling loops made from 23 gauge stainless steel tubing bent to form a loop shape approximately the size of primary auditory cortex. At the base of the loop, a micro-thermocouple made from twisting together PFA insulated copper (30 AWG; 0.254 mm) and constantan wire (Omega Engineering Limited, Manchester, UK), was soldered and secured with araldite. Thermocouple wires were soldered to a miniature thermocouple connector (RS components Ltd, UK) and secured with araldite.

Loops were surgically implanted over the border between middle and posterior ectosylvian gyrus, corresponding to the boundary between primary and non-primary auditory cortex. Surgery was performed in sterile conditions under general anesthesia, induced by a single intramuscular injection of medetomidine (Domitor; 0.1 mg kg-1; Orion, Finland) and ketamine (Ketaset; 5 mg kg-1; Fort Dodge Animal Health, Kent, UK). Animals were intubated and ventilated, and anesthesia was then maintained with 1.5% isoflurane in oxygen throughout the surgery. An i.v. line was inserted and animals were provided with surgical saline (9 mg kg-1) intravenously, while vital signs (body temperature, end-tidal CO2 and electrocardiogram) were monitored throughout surgery. General anesthesia was supplemented with local analgesic (Marcaine, 2 mg kg-1, Astra Zeneca) injected at the point of midline incision. Under anaesthesia, the temporal muscle overlying the skull was retracted and a craniotomy was made over the ectosylvian gyrus. The dura overlying the gyrus was retracted to allow placement of the cooling loop on the surface of the brain. The loop was shaped during surgery to best fit the curvature of the cortical surface prior to placement, and was then embedded within silicone elastomer (Kwik-Sil, World Precision Instruments) around the craniotomy, and dental cement (Palacos R+G, Heraeus) on the subject’s head. Bone screws (stainless steel, 19010-100, Interfocus) were also placed along the midline and rear of the skull (two per hemisphere) to anchor the implant. Implant anchorage was also facilitated by cleaning the skull with citric acid (0.1 g in 10 ml distilled water) and application of dental adhesive (Supra-Bond C&B, Sun Medical). Some temporal muscle and skin were then removed in order to close the remaining muscle and skin smoothly around the edges of the implant. Pre-operative, peri-operative and post-operative analgesia and anti-inflammatory drugs were provided to animals under veterinary advice.

Animals were allowed to recover for at least one month before resuming behavioral testing and beginning cortical inactivation experiments. To reduce the temperature of the cortical tissue surrounding the loop, cooled ethanol (100%) was passed through the tube using an FMI QV drive pump (Fluid Metering, Inc., NY, USA) controlled by a variable speed controller (V300, Fluid Metering, Inc., NY, USA). Ethanol was carried to and from the loop on the animal’s head via FEP and PTFE tubing (Adtech Polymer Engineering Ltd, UK) insulated with silicon tubing and, where necessary, bridged using two-way connectors (Diba Fluid Intelligence, Cambridge, UK). Ethanol was cooled by passing through a 1 meter coil of PTFE tubing held within a Dewar flask (Nalgene 4150±1000, NY, USA) containing dry ice and ethanol. After passing through the loop to cool the brain, ethanol was returned to a reservoir that was open to the air.

For a cooling session, the apparatus was first ‘pre-cooled’ before connecting an animal by pumping ethanol through spare cooling loops (i.e. loops that were not implanted in an animal) until loop temperatures fell below 0°C. The animal was then connected to the system, using the implanted thermocouples to monitor loop temperature at the cortical surface. The temperature was monitored online using a wireless transfer system (UWTC-1, Omega Engineering Ltd., Manchester, UK) or wired thermometer, and pump flow rates adjusted to control loop temperature.

For F1311, the animal was connected to the system and cooling began before the behavioral session, with the subject held by the experimenter and rewarded with animal treats (Nutriplus gel, Virbac, UK) while cooling pumps were turned on and loop temperatures reduced over five to ten minutes. When loop temperatures reached ≤12°C, the animal was placed in the behavioral arena and testing began. Ferret F1509 would not perform the task after being rewarded by the experimenter and so behavioral sessions were started and cortical temperature slowly reduced during task performance. Trials performed by the animal before the loops were cooled (≤20°C) were excluded from the analysis. For both animals, cooling took place while the animals were free to move without interaction with the experimenter and within the same apparatus as used for previous behavioral testing.

For both animals, the behavioral task during cooling was unchanged described above; i.e. the same ranges of sound levels were used and correction trials were included. For each trial in the task, the time of stimulus onset was recorded and cross-referenced with temperature records so that any trials in which cortical temperature was above threshold during a cooling session could be removed offline. As with all behavior, tests of vowel discrimination in continuous noise were performed on separate sessions from tests of discrimination in clean conditions and restricted noise. During testing, the experimenter was blind to the noise condition used on each trial.

To analyse the effects of cooling, we compared behavioral performance of each animal across multiple sessions: For one ferret (F1509), we compared the effects of cooling on paired testing sessions, performed on the same day. This was not possible for the other ferret (F1311) as cooled and control tests were not made on paired sessions. Therefore we compared behavior on cooled sessions with control sessions tested over the same period of time as cooling, but not necessarily on the same day. The effect of cooling was defined as the change in percent correct between cooled and control sessions (cooled – control). To test whether changes in performance were significant, we randomly shuffled the labels (cooled or control) for each trial and recalculated percentage correct values. This procedure was repeated 10^4^ times to generate the permutation distribution, from which we report the probability of observing a larger decrease in performance by chance. Significance of the permutation test was assessed at α = 0.05.

### Neural recording in behavior

Techniques for surgical implantation of electrodes, recording of neural activity and confirmation of electrode position have been described elsewhere (Town, Brimijoin et al. 2017). Briefly, each ferret was chronically implanted with two WARP microdrives (Neuralynx, MT), each containing sixteen independently moveable tungsten microelectrodes (WPI Inc., FL), placed over left and right auditory cortex. Neural activity was recorded continuously throughout a behavioral test session. Surgical procedures were the same as described for implantation of cooling loops, however the dura was not retracted prior to microdrive implantation. The bone screws used to anchor the implant were also used as ground and reference contact points, connected to each microdrive using tinned copper wire. Following implantation, animals were allowed to recover for at least one week before electrodes were initially descended, and a further two weeks before resuming behavioral testing.

On each electrode, voltage traces were recorded using TDT System III hardware (RX8 and RZ2) and OpenEx software (Tucker-Davis Technologies, Alachua, FL) with a sample rate of 25 kHz. To extract action potentials, data were band-pass filtered between 300 and 5000 Hz and motion artefacts were removed using a decorrelation procedure applied to all voltage traces recorded from the same microdrive in a given session (Musial, Baker et al. 2002). For each channel within the array, we identified candidate events as those with amplitudes between −2.5 and −6 times the RMS value of the voltage trace and defined waveforms of events using a 32-sample window centered on threshold crossings. Here we did not isolate single units by spike sorting, and considered multi-unit activity recorded at each site within cortex.

### Neural data analysis – Individual unit decoding

We identified 277 units that were responsive to sounds when data were combined across clean and restricted noise conditions. Responsiveness was defined as a significant change in firing rate measured in the 300 ms after onset of first vowel token from spontaneous activity in the 300 ms before making contact with the spout (Sign-rank test, p < 0.05). Responsiveness was initially assessed across all trials; however later analysis tested if units were responsive to sounds in clean or noise conditions separately. The same approach was taken to identify 128 units that were responsive to sounds when data were combined across clean, restricted and continuous noise conditions. Responsiveness was then tested separately for each experimental condition.

For each sound responsive unit, we decoded vowel identity (/u/ or /ε/) from single trial responses to sounds in a specific condition (e.g. clean sounds). The decoder used a simple spike-distance metric with leave-one-out cross-validation (LOCV) that is described in detail elsewhere (Town, Wood et al. 2018). Briefly, for a given dataset (e.g. 100 trials, with 50 trials per vowel) we removed a single test trial and calculated template responses for each vowel as the mean PSTH of responses on all other trials. We then assigned an estimate of the vowel on the test trial as the template with the smallest Euclidean distance to the test trial. Where equal distances were observed between test trial and multiple templates, we randomly estimated (i.e. guessed) which of the equidistant templates was the true stimulus feature. This procedure was repeated for all trials and decoding performance was measured as the percentage of trials on which the vowel was correctly recovered. To ensure adequate data for template construction, we only considered units with responses tested on a total of ≥ 10 trials. In addition to decoding vowel identity, we also decoded the behavioral response on each trial (go left or right) using the same procedure. Both correct and error trials were included in each decoding analysis.

Neurons in auditory cortex have a wide variety of response profiles that makes it difficult to select a single fixed time window over which to decode neural activity. To accommodate the heterogeneity of auditory cortical neurons and identify the time at which stimulus information arose, we repeated our decoding procedure using different time windows varying in start time (−0.5 to 1 s after stimulus onset, varied at 0.1 s intervals) and duration (10 to 500 ms, 10 ms intervals, yielding a total of 1550 possible windows). Within this space, we then reported the parameters that gave best decoding performance, and where several parameters gave best performance, we reported the time window with earliest start time and shortest duration.

To assess the significance of decoding performance, we conducted a permutation test in which the decoding procedure (including the full temporal optimization) was repeated 100 times but with the decoded feature randomly shuffled between trials to give a null distribution of decoder performance. The null distribution of shuffled decoding performance was then parameterized by fitting a Gaussian probability density function, which we subsequently used to calculate the probability of observing the real decoding performance. Units were identified as informative when the probability of observing the real performance after shuffling was less than 5%.

In Supplementary Figure 6, we ran an additional decoding analysis to test for robust representations of vowel identity across clean and restricted noise conditions. Here, we built templates (trained the decoder) from data in a specific condition (e.g. using responses to clean sounds) and then tested decoding performance on data from a different condition (e.g. responses in noise). The decoder used the same temporal optimization and permutation test procedure, but indicated if information about vowel identity was conserved across different stimulus sets (i.e. noise regimes). We repeated the decoding in both directions: training in clean conditions and testing responses to sounds in noise, or alternatively training on noise data and testing on clean data. The same procedure was then repeated when labels on test data were randomly shuffled and decoding performance recomputed. Shuffles were repeated 1000 times in order to identify units that performed significantly better than chance (permutation test, p < 0.05). The analysis was only conducted for comparisons of responses to sounds in clean conditions or temporally restricted noise.

### Neural data analysis – Population decoding

To decode vowel identity from the single trial responses of populations of units, we summed the number of units that estimated each stimulus, weighted by the confidence of each unit’s estimate, and took the stimulus with the maximum value as the population estimate on that trial. Confidence weights for individual unit (*w*) estimates were calculated as:

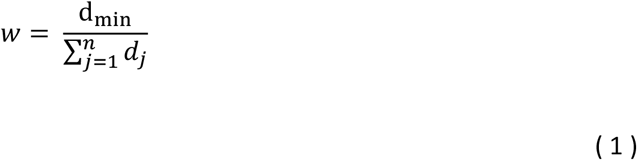

Where *n* was the number of vowels and *d* was the spike distance between the test trial response and response templates generated for each stimulus class. Here, d_min_ represents the minimum spike distance that corresponded to the estimated stimulus for that unit.

We tested populations of 200 units for which we recorded neural responses on at least 8 trials for each vowel. Within this subpopulation, we randomly sampled one hundred different populations, each with a unique combination of 200 units, without replacement from the whole population of sound-responsive units available (n = 277).

### Frequency tuning

We assessed frequency tuning of units by presenting pure tones to animals listening passively while receiving water at the center spout in the same behavioral chamber in which vowel discrimination was tested. For each unit tested, we compared sound-evoked responses to pre-stimulus activity to filter out units that were not responsive to pure tones. For 89 tone-responsive units, we then plotted the frequency response area and measured the best frequency as the frequency that generated the strongest mean firing rate.

### Neural recording during cooling

Recordings in suprasylvian cortex (SSY) during cooling were performed under anesthesia induced with medetomidine (Domitor; 0.022mg/kg/h; Pfizer, Sandwich, UK) and ketamine (Ketaset; 5mg/kg/h; Fort Dodge Animal Health, Southampton, UK), and maintained by intravenous infusion of (5 ml/h) of a mixture of medetomidine and ketamine in lactated Ringers solution augmented with 5% glucose, atropine sulfate (0.06 mg/kg/h; C-Vet Veterinary Products) and dexamethasone (0.5 mg/kg/h, Dexadreson; Intervet, UK). The animal was intubated and placed within a stereotaxic frame, after which the left temporal muscle was largely removed in order to perform a craniotomy that exposed the suprasylvian and pseudosylvian sulci. The dura was removed over SSY and the brain protected with 3% agar solution. The eyes were protected with zero-refractive power contact lenses. The animal was then transferred to a sound-attenuating chamber, where body temperature, end-tidal CO_2_, and the electrocardiogram were monitored throughout recording.

Neural activity was recorded with 16-channel, single shank silicon probe (A1×16, Neuronexus Technologies, Ann Arbor, MI) placed at the center of a cooling loop, which was itself placed on the suprasylvian cortex, in the region posterior to auditory cortex (See Atilgan, Town et al. 2018 for further details). The design of the cooling loop system was similar to that designed for behaving experiments, with the exception that the loop was not embedded within dental cement during anesthetized experiments. Following loop placement on the cortical surface, the silicon probe was oriented perpendicular to the cortical surface and descended until spiking activity was detected on the majority of channels. Several electrode penetrations within the vicinity of the loop were made for each ferret, but the loop remained in a fixed position across penetrations. Additional recordings were performed outside the vicinity of the cooling loop, in ectosylvian gyrus (auditory cortex), but are not considered here.

Following a period where the electrode was left alone to ensure stability of recording (5 – 30 minutes), ferrets were presented with 100 ms stimuli that were either auditory (broadband noise), visual (white LED) and audiovisual stimuli (noise + LED)(Atilgan, Town et al. 2018). Each stimulus was repeated 20 – 40 times in a random order with a jittered interval between presentations (minimum: 900 ms). Neurons in suprasylvian cortex were only driven by visual stimuli and thus we only used visual responses to model the effects of cooling. Tests of cooling began with identification of sites in which visual responses were observed while the cortex was at its normal temperature; the pump was then activated, driving cooled ethanol through the system, and tests of neural responsiveness were repeated at 5 – 10 minute intervals as the cortical temperature decreased to between 5 and 10°C (Wood, Town et al. 2017). Cooling was then suspended and the cortical temperature allowed to recover to body temperature.

### Simulated cortical inactivation

From recordings in SSY, we identified 83 units that were responsive to visual stimuli using a sign-rank test comparing pre- and post-stimulus activity in 150 ms windows. Responsiveness was assessed using control data from responses before cooling. We then compared firing rates in control and cooled conditions in 10 ms bins, from 100 ms before, to 800 ms after stimulus presentation. A linear regression model was fitted to quantify the association between firing rates in control (*x*) and cooled conditions (*y*):

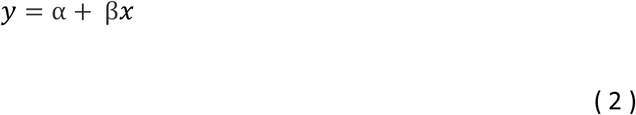

We identified significant model fits for 73 / 83 units, and plotted the gradient of the regression line (β) against the cortical temperature achieved during cooling. The gradient served as a proxy measure for the suppression of neural activity from which we could estimate the proportion of spikes suppressed for a given cortical temperature. To simulate the effects of cooling in vowel discrimination, we then randomly deleted a proportion of spikes from responses of auditory cortical neurons recorded during task performance described above. The percentage of spikes deleted was a key parameter in our simulations, as was the proportion of neurons that underwent spike deletion (i.e. that were ‘cooled’). For a given percentage of deleted spikes and affected neurons, we reran our population decoder using the same approach described above to recover vowel identity from neural activity. Here we only used neural activity in a 100 ms window following stimulus onset. We simulated different effects of cooling by varying the percentage of deleted spikes (0 to 100% in 10% steps) and proportion of affected units (0 to 100% in 10% steps).

## Acknowledgements

This work was funded by a Royal Society Dorothy Hodgkin Fellowship to JKB, the BBSRC (BB/H016813/1) and a Wellcome Trust / Royal Society (WT098418MA). We are grateful to Bhavisha Parmar for discussions on the role of noise-reduction technologies in hearing aids and cochlea implants.

## Author Contributions

SMT and JKB wrote the paper; all authors collected the data; SMT analyzed the data

---

**Supplementary Figure 1.**
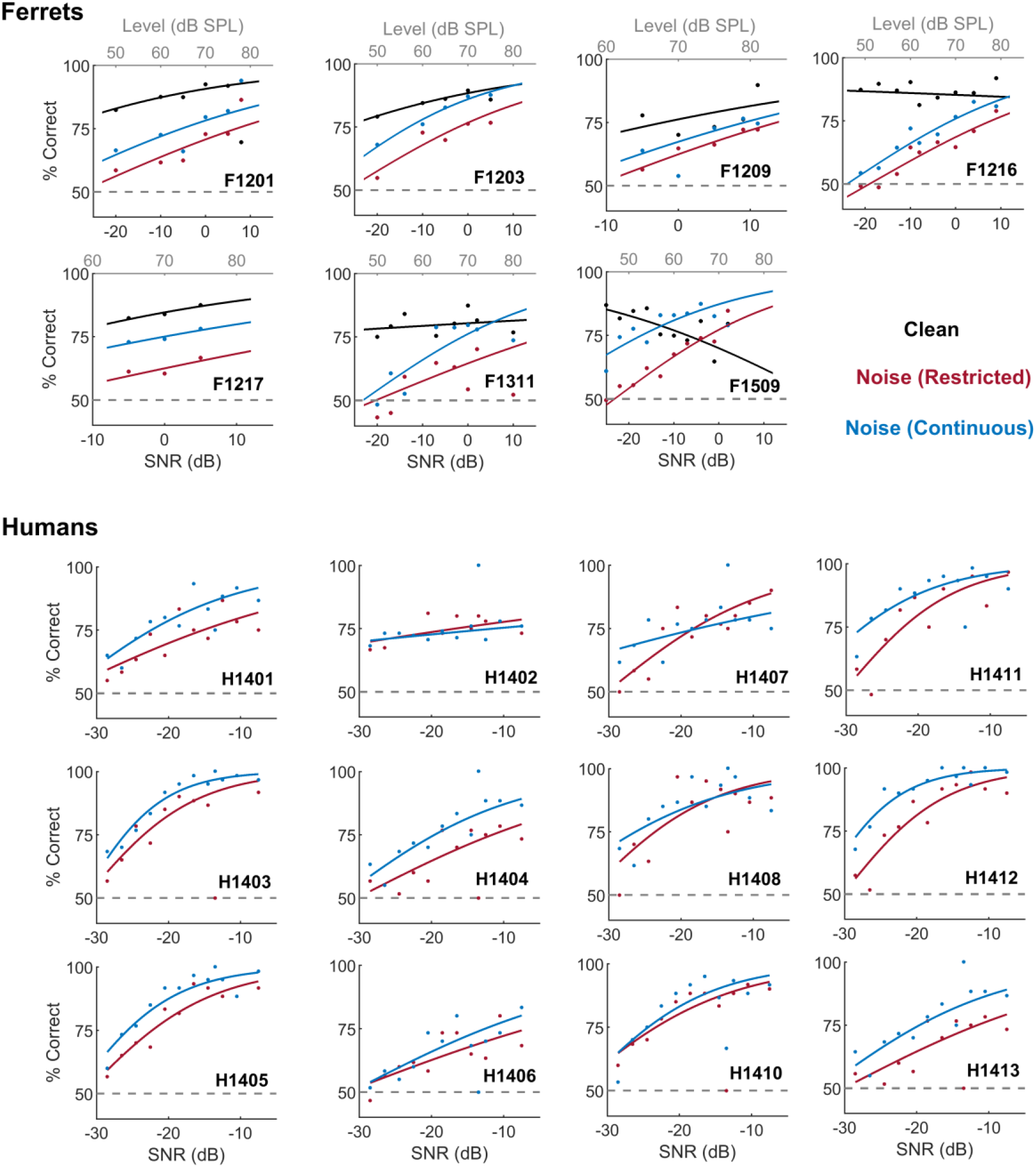
Task performance for all subjects. Performance of all ferrets and humans tested with vowels in clean and noise conditions. Data points show performance at specific signal levels or SNRs; filled lines showing logistic regression fits to data. Dashed grey in each plot indicates chance performance (50%).

**Supplementary Figure 2.**
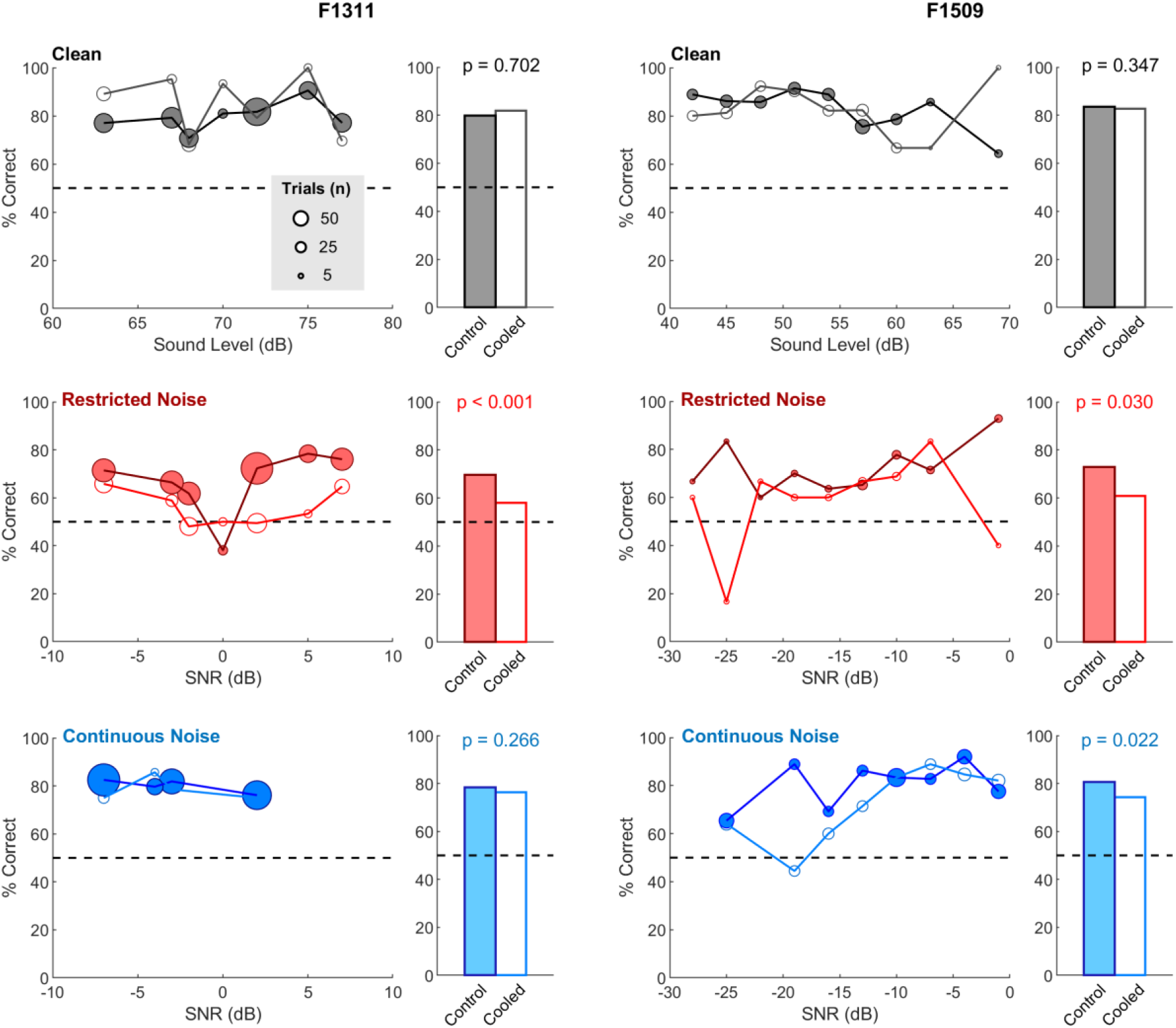
Effects of cooling by sound level and SNR. Vowel discrimination performance of two ferrets (F1311 and F1509) in clean and noise conditions during cooling and control testing. P values indicate effect of cooling across signal levels or SNRs (permutation test, observed deficit vs. 10^4^ iterations).

**Supplementary Figure 3.**
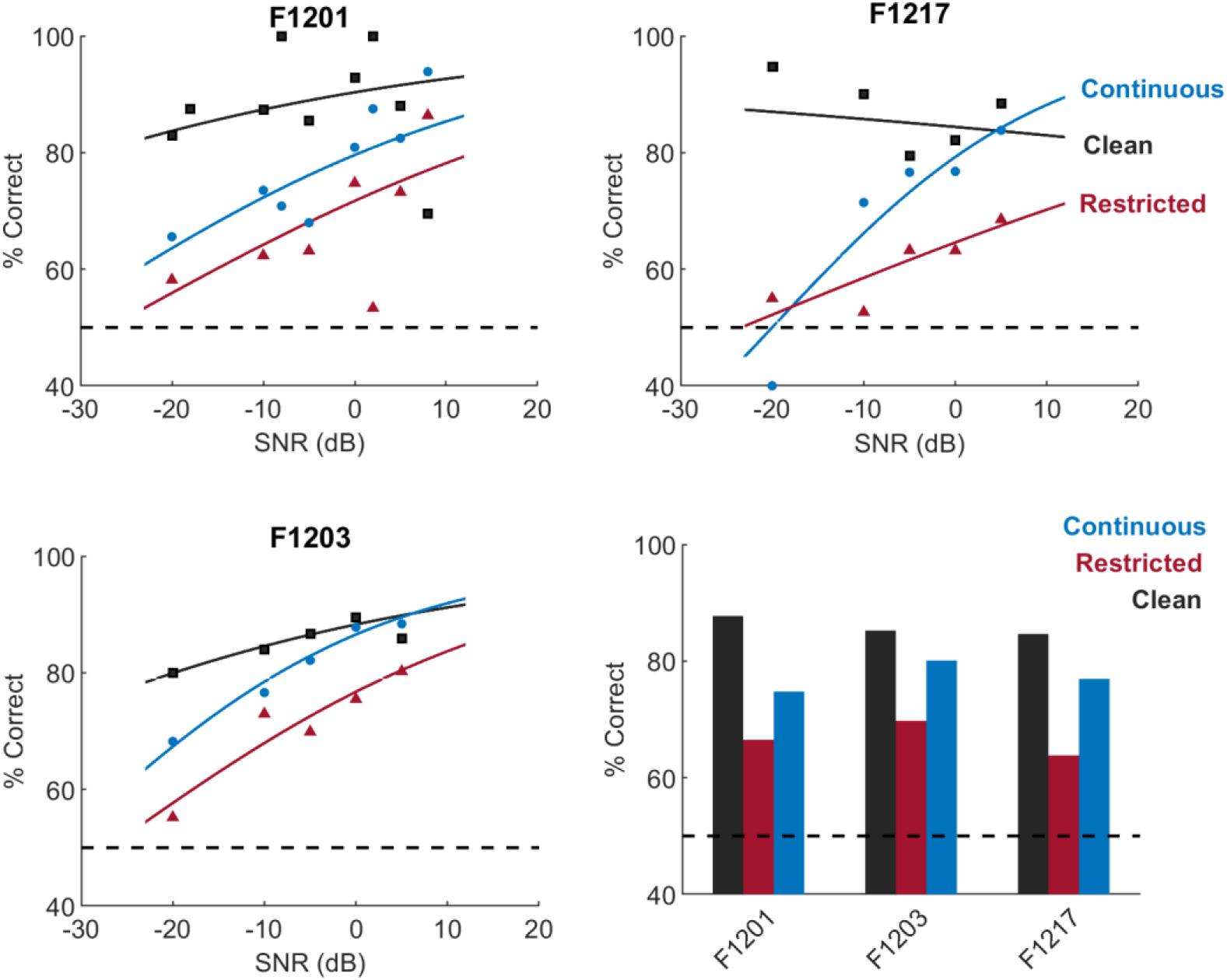
Behavior of each ferret during neural recording. Scatter plots show performance at each SNR value tested in clean (black squares), continuous (blue circles) or restricted (red triangles) noise conditions. Full lines show logistic regression models fitted separately for each noise type; dashed black lines indicate chance performance. Bar plots show performance across all SNRs for each ferret.

**Supplementary Figure 4.**
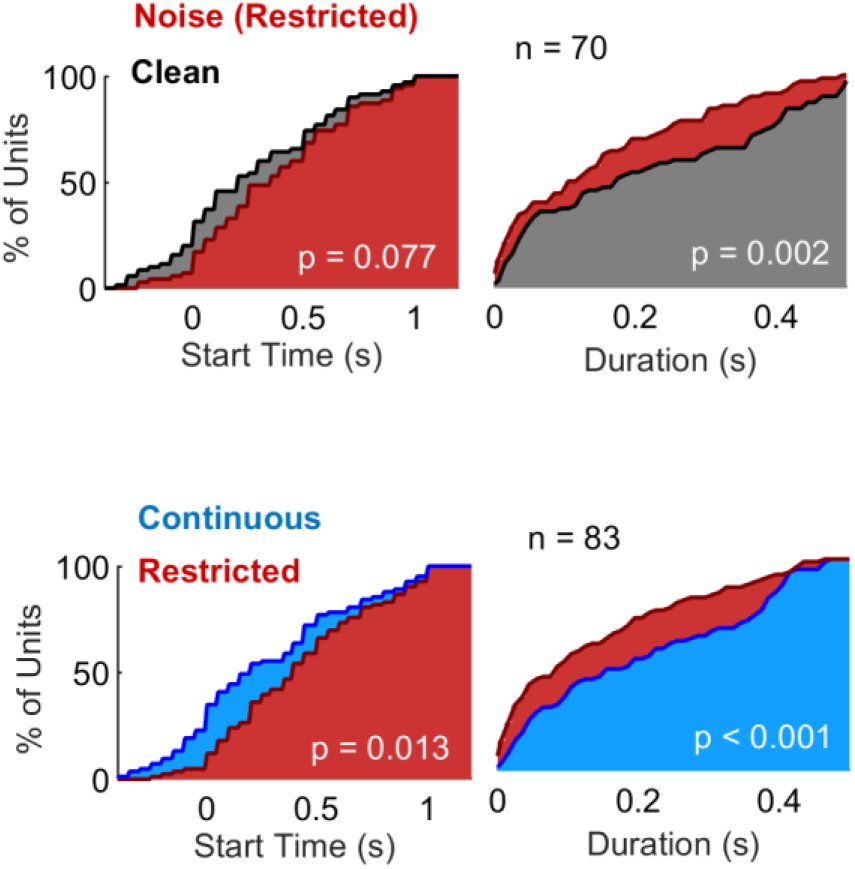
Effects of noise on timing parameters in optimized vowel decoding. Start time and duration of time windows that gave best decoding performance following optimization. The time window required to decode vowel identity ‘optimally’ was significantly shorter in temporally restricted noise than clean conditions (sign-rank comparison of window duration: time difference (Δt) = 50 ms, p = 0.002). Decoding also began later in noise than clean conditions, although this difference was not significant (Δt: 175 ms, sign-rank test, p = 0.077). Data shown as cumulative distribution of units that were informative about vowel identity in clean and/or noise conditions (n = 70). Optimized decoding of vowel identity in continuous noise began significantly earlier (sign-rank test, median difference = 100 ms, p = 0.013) and lasted significantly longer (median difference = 50 ms, p < 0.001) than in restricted noise. Data shown for units that were informative about vowel identity in clean, restricted and/or continuous noise conditions (n = 83).

**Supplementary Figure 5.**
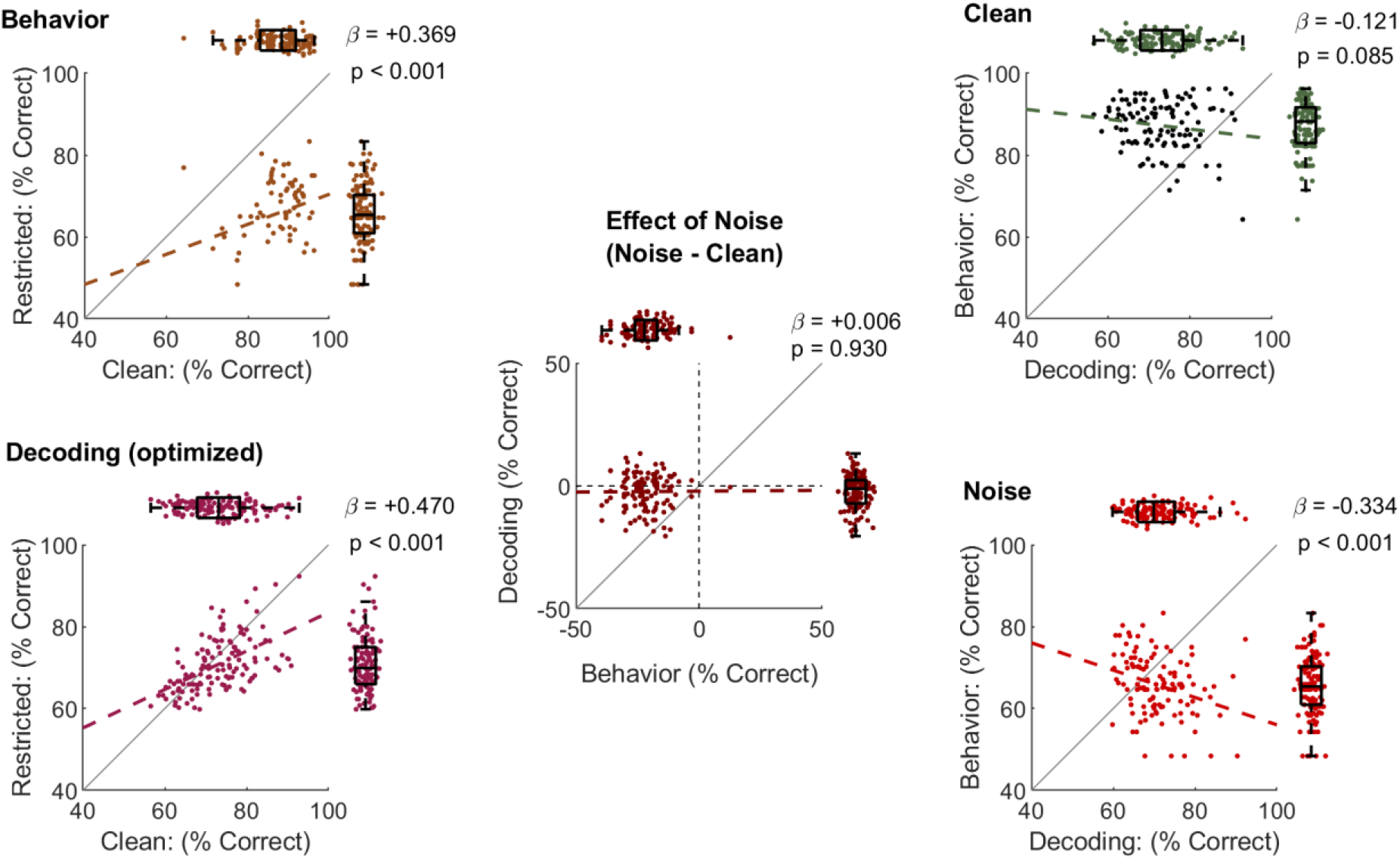
Correlations between decoding performance and behavior. Comparison of behavioral and decoding performance in clean and restricted noise for units that were responsive to sounds in both conditions (n = 130). Behavior shows ferrets’ task performance across all trials for which data from each unit was recorded. Decoding (Optimized) shows performance estimating vowel identity from single trial neural responses (see also Fig. 2B). Effect of noise shows difference in decoding and behavioral performance. Clean and noise plots show behavioral and decoding performance for each unit. Boxplots show median, interquartile range and 99.3% confidence interval for each dimension. Values (p, β) show the regression coefficient and significance of correlation between variables for each plot.

**Supplementary Figure 6.**
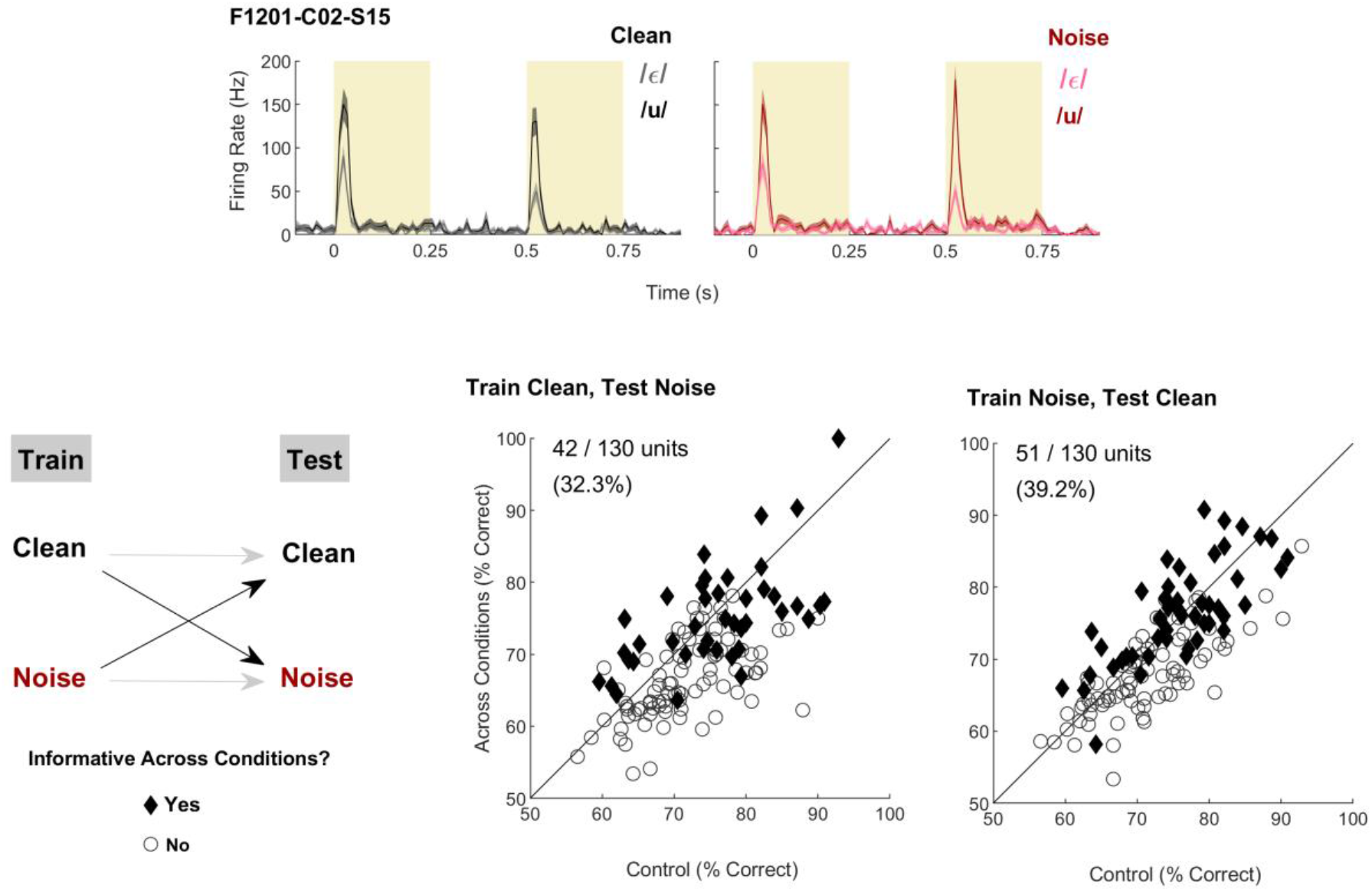
Noise-tolerant coding in Auditory Cortex. Top: Example responses from one unit that was informative about vowel identity in both clean and noise conditions. This unit was representative of a subpopulation of units (26 / 130; 20%) for which vowel identity could be successfully decoded across conditions when using independently optimised decoders (see also Fig. 2B). Bottom: Performance decoding vowel identity across conditions either training on responses to clean sounds and testing responses to sound in noise (left) or vice versa (right). In both cases, performance is shown vs. decoding of clean sounds when trained using responses to clean sounds. Values show the proportion and percentage of units for which test decoding performance was significant, indicating conserved representation of vowel identity across noise conditions. (permutation test, n = 10^3^ iterations, *p* < 0.05).

**Supplementary Table 1.**
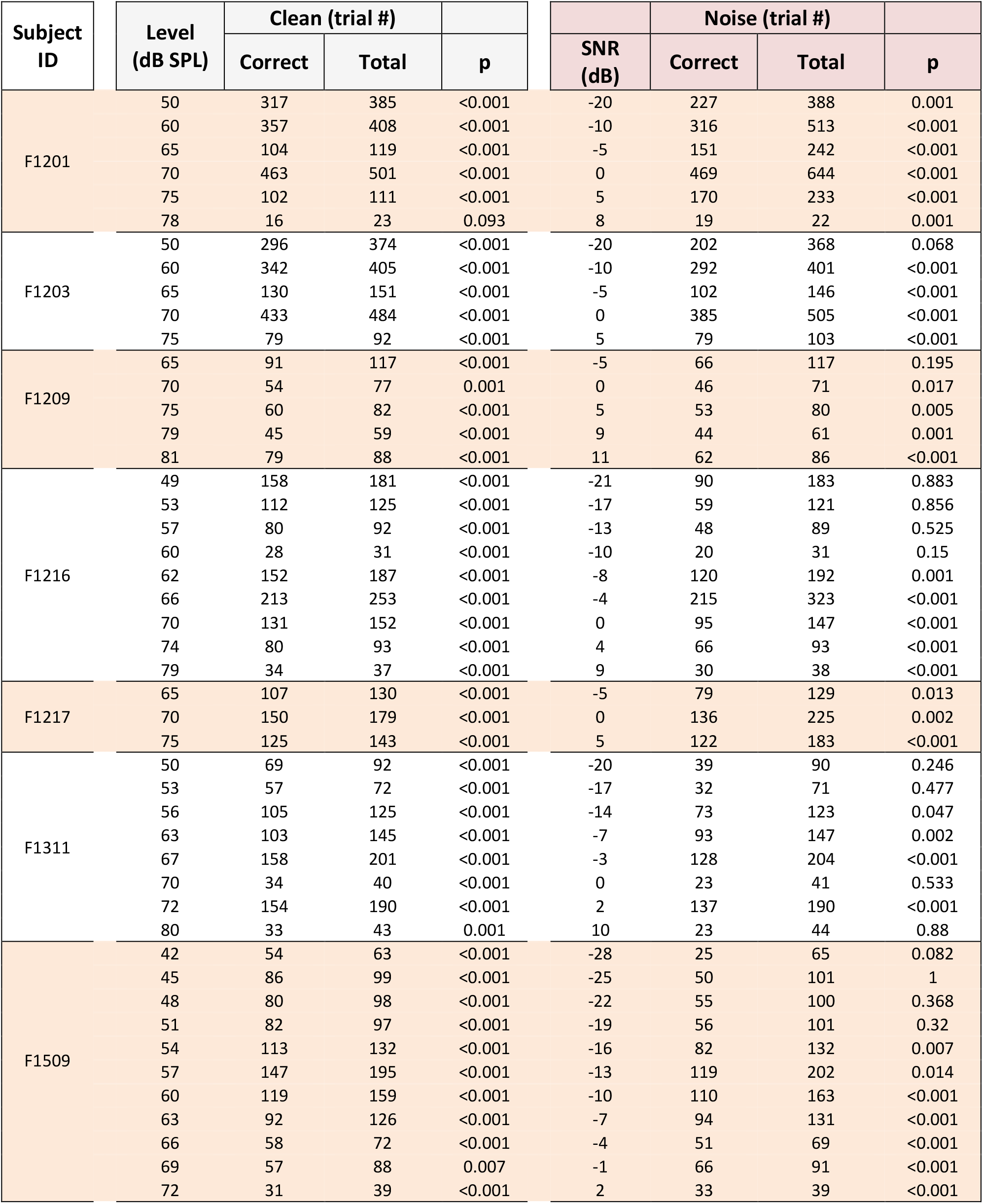
P values show uncorrected probability of the observed performance resulting from binomial test.

**Supplementary Table 2.** Coefficients resulting from logistic regression between sound level or SNR and proportion of trials correct. Positive coefficients indicate that performance increased with sound level or SNR. For two animals we observed negative coefficients (blue) indicating performance decreased with increasing sound level in clean conditions.

| Subject ID | Clean |  | Noise |  |
| --- | --- | --- | --- | --- |
|  | Coefficient | p | Coefficient | p |
| F1201 | 0.035 | < 0.001 | 0.032 | < 0.001 |
| F1203 | 0.034 | < 0.001 | 0.044 | < 0.001 |
| F1209 | 0.032 | 0.0934 | 0.043 | 0.010 |
| F1216 | -0.006 | 0.567 | 0.041 | < 0.001 |
| F1217 | 0.039 | 0.241 | 0.025 | 0.2821 |
| F1311 | 0.004 | 0.704 | 0.032 | 0.001 |
| F1509 | -0.036 | < 0.001 | 0.053 | < 0.001 |

**Supplementary Table 3.** Regression analyses comparing behavioral and decoding performance

| Time | Units | Predictor | Response | Coefficient Estimate | P |
| --- | --- | --- | --- | --- | --- |
| Optimized | All<br>(n = 130) | Clean Decoding | Clean Behavior | -0.121 | 0.0847 |
|  |  | Noise Decoding | Noise Behavior | -0.333 | 0.0004 |
|  |  | Clean Behavior | Noise Behavior | 0.369 | 0.0002 |
| | | Clean Decoding | Noise Decoding | 0.470 | $1.2 \times 10^{-11}$ |
|  |  | Noise Effect on Behavior | Noise Effect on Decoding | 0.007 | 0.929 |
|  | Informative<br>(n = 70) | Clean Decoding | Clean Behavior | -0.140 | 0.157 |
|  |  | Noise Decoding | Noise Behavior | -0.254 | 0.0278 |
|  |  | Clean Behavior | Noise Behavior | 0.229 | 0.0782 |
|  |  | Clean Decoding | Noise Decoding | 0.330 | 0.002 |
|  |  | Noise Effect on Behavior | Noise Effect on Decoding | -0.0519 | 0.674 |
| Fixed (0 – 100 ms) | All<br>(n = 130) | Clean Decoding | Clean Behavior | 0.0160 | 0.815 |
|  |  | Noise Decoding | Noise Behavior | -0.0471 | 0.354 |
|  |  | Clean Decoding | Noise Decoding | 0.2508 | 0.0009 |
|  |  | Noise Effect on Behavior | Noise Effect on Decoding | -0.0527 | 0.697 |
|  | Informative<br>(n = 70) | Clean Decoding | Clean Behavior | -0.071 | 0.332 |
|  |  | Noise Decoding | Noise Behavior | -0.0591 | 0.542 |
|  |  | Clean Decoding | Noise Decoding | 0.115 | 0.246 |
|  |  | Noise Effect on Behavior | Noise Effect on Decoding | -0.0104 | 0.955 |

**Supplementary Table 4.** Binomial test results used to determine if vowel discrimination was significantly greater than chance performance. Performance did not significantly exceed chance levels in only one condition (restricted noise) in one animal (F1509, 48/79 trials correct [60.8%], Binomial test: p = 0.071), although this most likely reflected the smaller sample sizes obtained during cooling rather than a total abolition of task performance.

| Subject | Condition |  | Trials (n) |  | Binomial Test |
| --- | --- | --- | --- | --- | --- |
|  |  |  | Correct | Total |  |
| F1311 | Clean | Control | 516 | 646 | $p < 0.001$ |
| | | Cooled | 172 | 210 | $p < 0.001$ |
| | Restricted | Control | 601 | 864 | $p < 0.001$ |
| | | Cooled | 238 | 411 | $p = 0.002$ |
| | Continuous | Control | 999 | 1274 | $p < 0.001$ |
| | | Cooled | 74 | 97 | $p < 0.001$ |
| F1509 | Clean | Control | 229 | 274 | $p < 0.001$ |
| | | Cooled | 172 | 208 | $p < 0.001$ |
| | Restricted | Control | 78 | 107 | $p < 0.001$ |
| | | Cooled | 48 | 79 | $p = 0.071$ |
| | Continuous | Control | 274 | 340 | $p < 0.001$ |
| | | Cooled | 219 | 295 | $p < 0.001$ |

